# ClimLimits: a global database of multivariate realized climate limits for animal species

**DOI:** 10.64898/2026.07.16.738937

**Authors:** Krishna S. Girish, Vasilis Dakos, Claire Jacquet

## Abstract

**Introduction and Aim:** Assessing the realized climate limits for a species based on the climate conditions (i.e., different aspects of temperature and precipitation) a species has experienced over its range enables us to determine the climatic boundaries of its existence, and thus its potential exposure to novel climate conditions in the future. We combine species’ range maps from IUCN and BirdLife International with global climate data from the ERA5 reanalysis and five Earth System Models (ESMs) to produce ClimLimits: a database of multivariate realized species climate limits based on historical temperature and precipitation for terrestrial and freshwater animal species worldwide.

**Main variables included:** For a total of 54,255 species (24,731 terrestrial, 18,182 freshwater, and 11,342 terrestrial-freshwater species), we estimated 44 species climate limits, which delineate the most extreme climate conditions experienced by a species over its entire range over the last 80 years (1940-2020). The database accounts for three aspects of species climate limits: a) maximum and minimum values of temperature and precipitation experienced over the historical reference period, b) maximum annual variability in temperature and precipitation, and c) maximum frequency, intensity, duration and severity of extreme events (heatwaves, cold-spells, and droughts). Climate data is sourced from the ERA5 reanalysis and from five different Earth System Models (ESMs), producing 6 different subsets of the ClimLimits database.

**Time coverage:** Species climate limits are estimated based on historical climate records from 1941-2014 (5 ESMs) and 1940-2020 (ERA5). Temperature-based limits are inferred at a daily scale, while precipitation-based limits are inferred at a monthly and yearly scale.

**Spatial coverage:** Global, over 24km x 24km grid-cells.

**Taxa:** Terrestrial and freshwater taxa, including amphibians, birds, mammals, reptiles, freshwater fish, and freshwater invertebrates, with shapefiles from IUCN and BirdLife International. Data is produced at the species level.

**Applications:** ClimLimits provides ready-to-use standardized realized climate limits for individual species across multiple aspects of climate, facilitating global-scale assessments of macroecological patterns and climate exposure risk for species.

## 1 Introduction

Projected future climate change is increasingly being understood as a major driver of biodiversity loss (Parmesan and Yohe, 2003; Urban, 2015; Malhi et al., 2020). Novel climate conditions can affect species in many ways, from forcing them to acclimate (Gunderson and Stillman, 2015; Weaving et al., 2022) or to move ranges to track their optimal environmental conditions (Lenoir et al., 2020), to affecting their behavior and physiology (Kingsolver et al., 2013; Sunday et al., 2014), and eventually inducing lethal conditions that cause species to become locally or globally extinct (Thomas et al., 2004; Wiens, 2016; Freeman et al., 2018). These climatic changes can occur across multiple climate variables at once, modifying many facets of local climate limits. For example, future climate in many parts of the world may become both hotter and drier (De Luca and Donat, 2023). Thus, concerted changes in many local climate limits could amplify extinction risk to biodiversity as species are simultaneously exposed to multiple physiologically stressful conditions (Halpern et al., 2015; Kefford et al., 2023).

While studies have looked into climate change exposure risk by modeling species’ realized niche limits focusing only on single climate limits (either historical maximum temperatures or extreme heat events (Trisos et al., 2020; Murali et al., 2023; Pigot et al., 2023), there is a growing interest in multivariate climate limits at various spatial scales (Pearson et al., 2006; Valavi et al., 2022; Karger et al., 2023; Adde et al., 2025). In addition to increases/decreases in global temperature and precipitation averages, climate change is understood to also modify other aspects of temperature and precipitation; each of these aspects have different global patterns and trends across ecosystems (Garcia et al., 2014). These aspects involve novel maxima/minima, changes in variability, and the occurrence of extreme events across different climate variables (i.e. temperature and precipitation). Since different species may have varying levels of sensitivity to changes in any of these aspects of climate conditions, computing climate limits across multiple aspects of climate variables can provide a more complete picture of climate change sensitivity across a species, taxon, or species assemblage. For example, in addition to increases in mean temperature, temperature variability is projected to increase in much of the tropics (Bathiany et al., 2018), alongside global increases in the frequency, duration and intensity of extreme heat and drought events (Gao et al., 2023). Therefore, there is an urgent need to synthesize species’ climate limits across multiple aspects of climate to better assess species’ risk to future climate change.

To support research into species’ multivariate climate limits for climate change risk assessment, we build the ClimLimits database. Using expert-curated range maps from the IUCN for 54,255 terrestrial and freshwater species sourced from 80,016 different populations, combined with fine-scale global reconstructions of historical climate (i.e. temperature and precipitation) conditions from the ERA5 reanalysis (from 1940-2020) (Hersbach et al., 2023) and 5 Earth System Models (ESMs) (from 1941-2014) (Lange, 2019), we compute realized local climate limits across the planet at the level of 24km x 24km grid-cells, and translate these local climate limits to species climate limits- that is, the minimum/maximum local climate limits experienced by species over their entire ranges. For each species, we annotate 44 different species climate limits (Table 1), encompassing 3 aspects of the realized climate conditions experienced by a species over its geographic range roughly from 1940 to 2020: a) maximum and minimum values of temperature and precipitation (expressed as percentiles), b) maximum annual variability in temperature and precipitation, and c) maximum frequency, intensity, duration and severity of extreme events (i.e., heatwaves, cold-spells, droughts). To account for model output differences between ERA5 reanalysis and the five ESMs, we compute the 44 species climate limits for each of the six historical climate records separately.

**Table 1:**
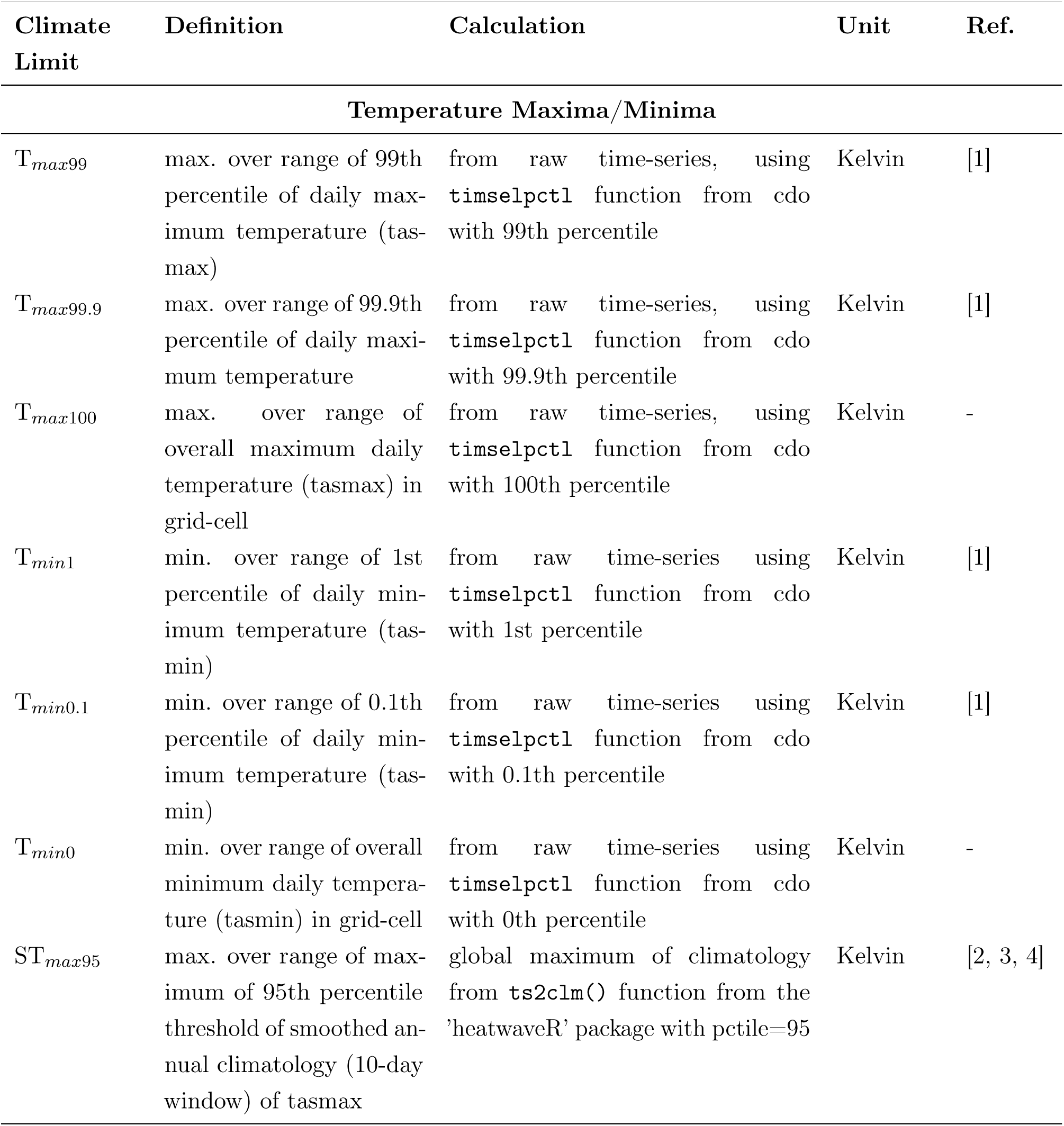
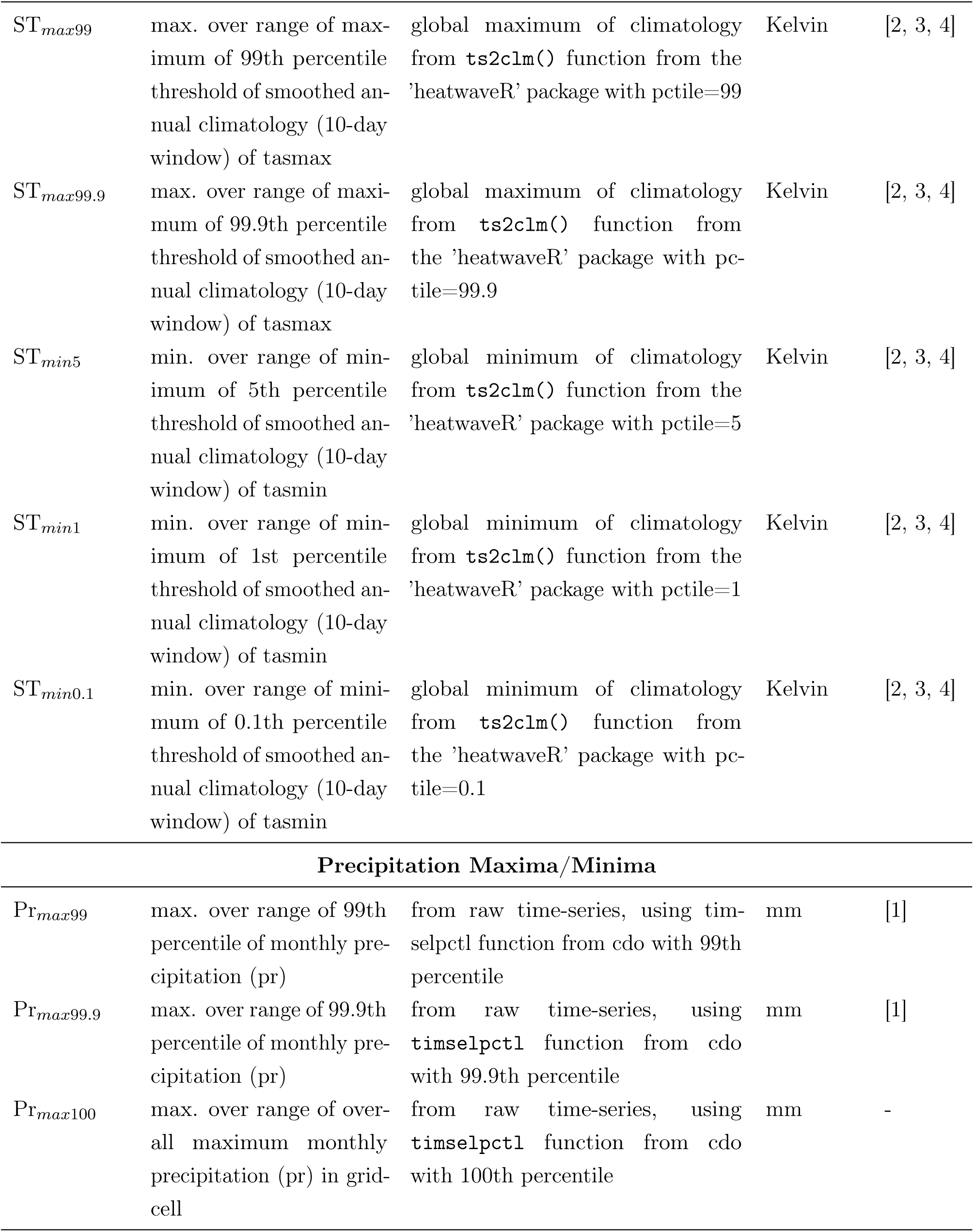
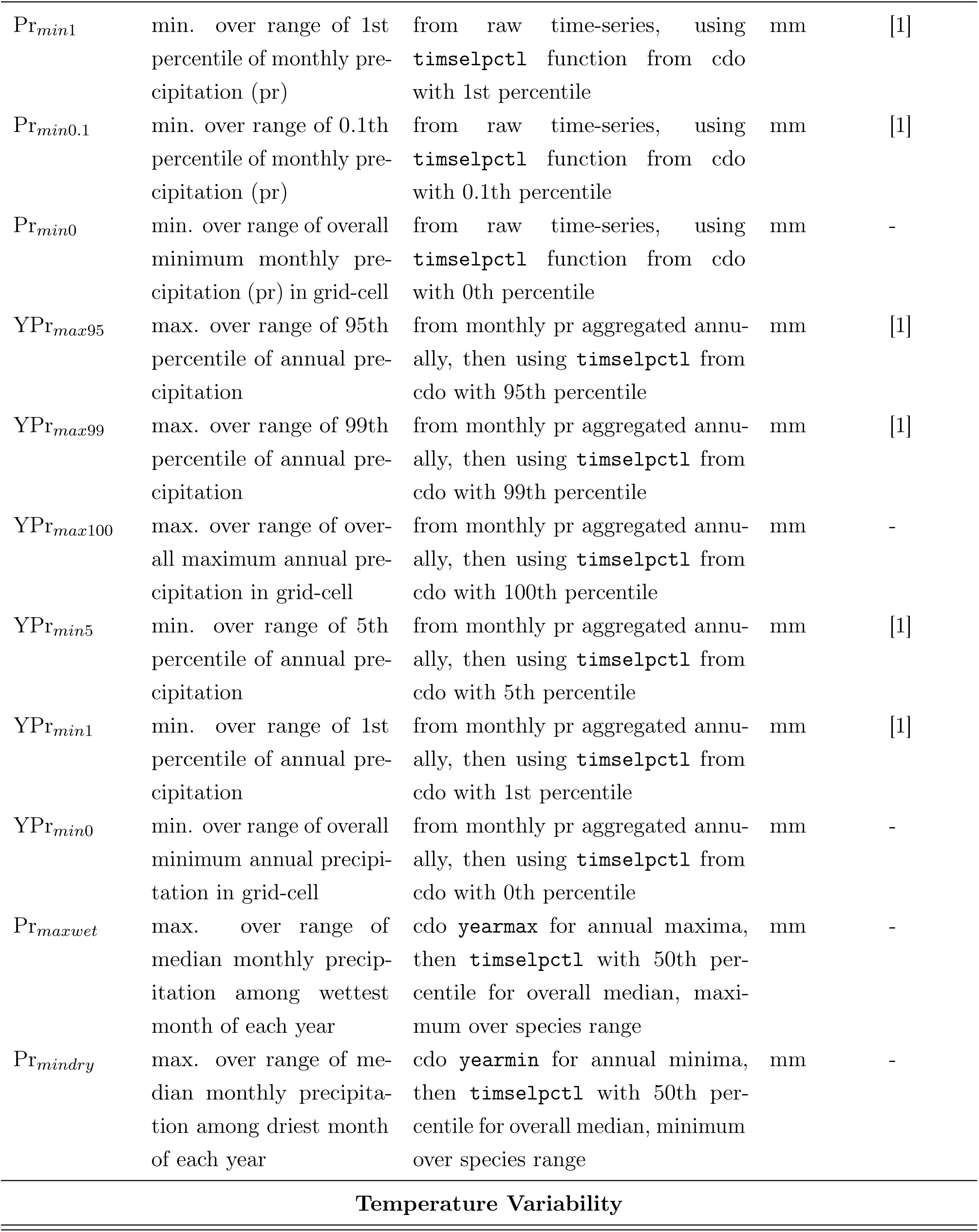
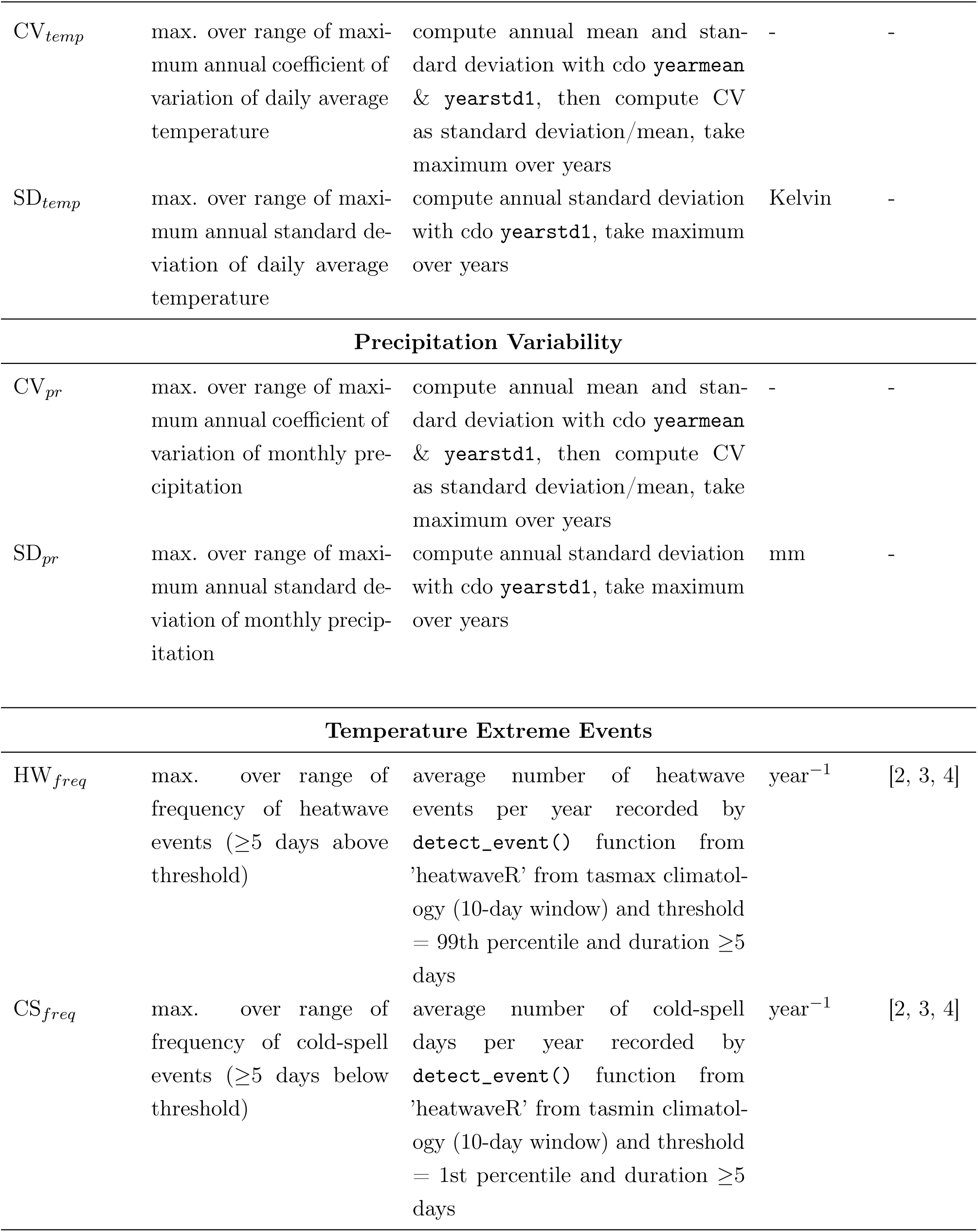
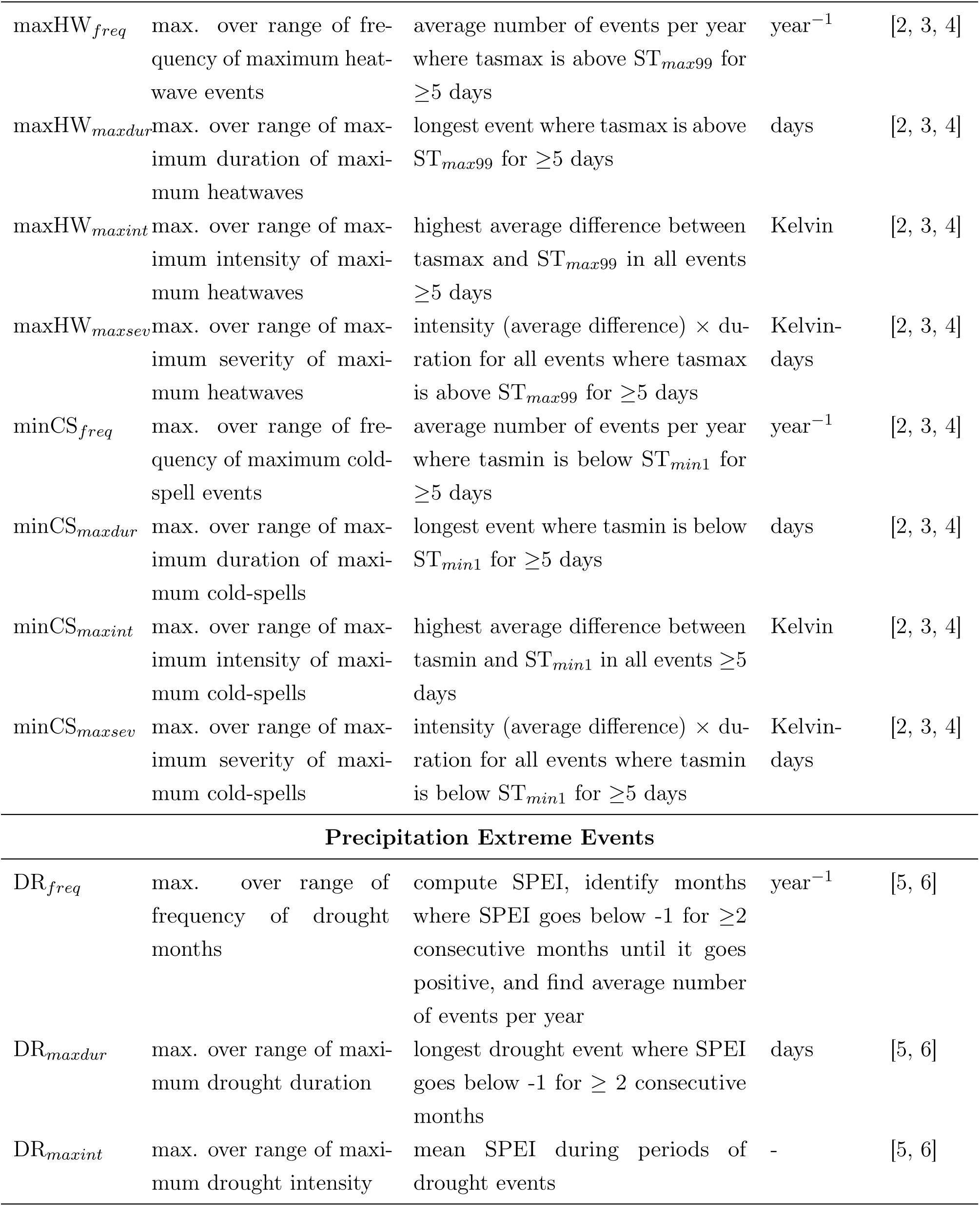
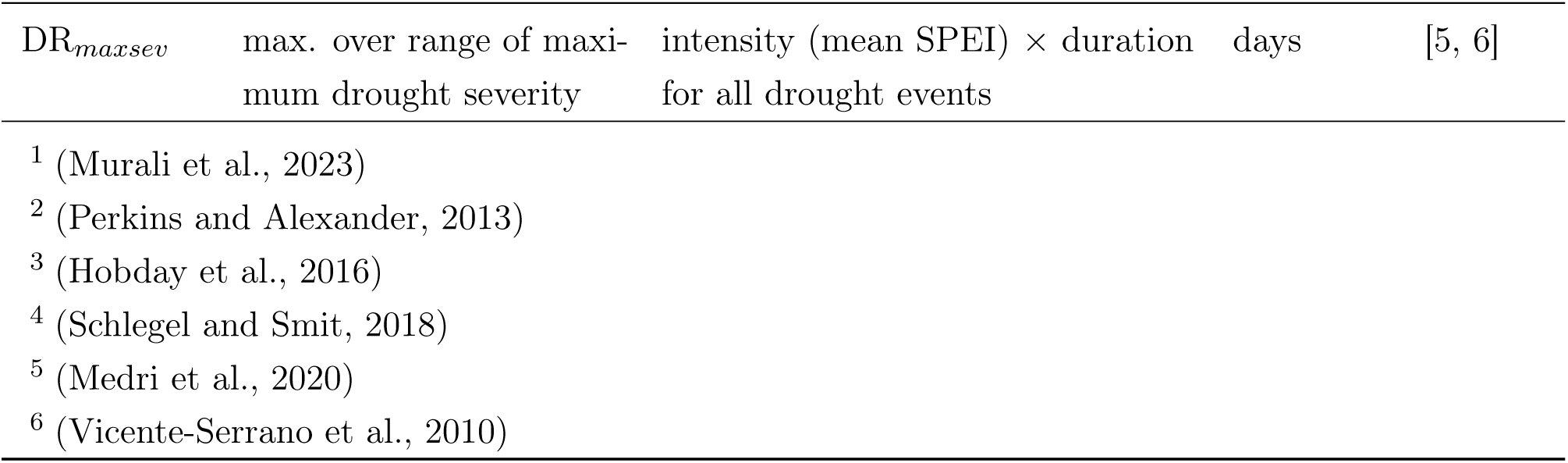
The 44 species climate limits (in temperature and precipitation), definitions, calculation, units in ClimLimits, alongside the primary reference of its estimation. Species climate limits are computed by taking the maximum/minimum of local climate limit values from each grid-cell in the species’ range. Climate limits are divided in six categories: Temperature Maxima/Minima, Precipitation Maxima/Minima, Temperature Variability, Precipitation Variability, Temperature Extreme Events, Precipitation Extreme Events.

Our database includes a wide range of simple, interpretable species climate limits with multiple variants that allow the quantification of realized climate limits in a wide range of settings, mitigating geographical and taxonomic biases that often plague macroecological biodiversity studies (Troudet et al., 2017; Trull et al., 2018). Built according to FAIR principles (Michener and Jones, 2012; Garnier et al., 2025), we anticipate the ClimLimits database to be useful for comparing macroecological trends across groups of species, assessing climate change vulnerability at the species/community/ecosystem level, exploring species extinction risk and adaptive capacity under future climate scenarios, and overall supporting environmental policy and conservation assessments.

## 2 Methods

The method for computing species climate limits is summarized for a sample species in Figure 1. In the following subsections, we detail how we estimated each local climate limit, and translated them into species climate limits (the full list is in Table 1).

**Figure 1:**
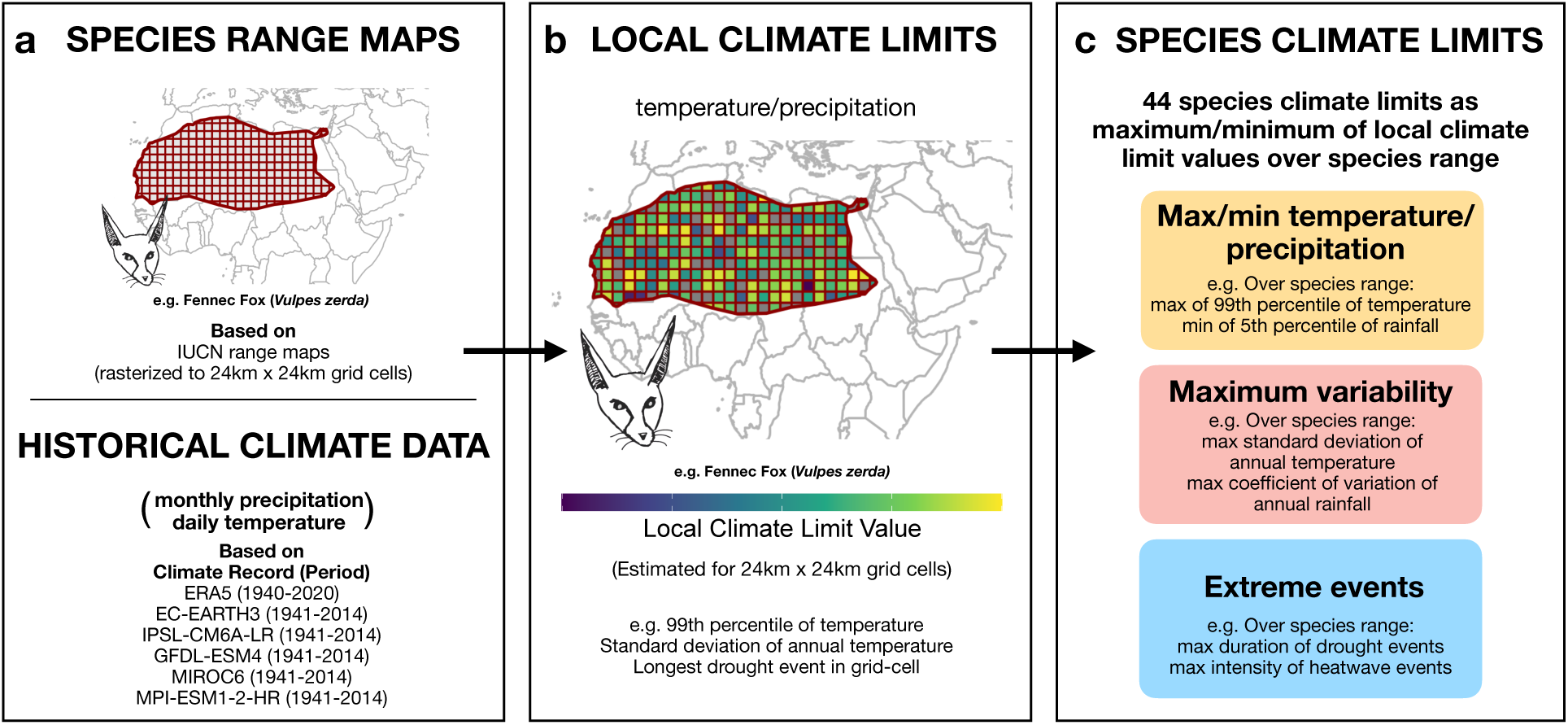
The building blocks of the ClimLimits database. Estimation of species climate limits from range and climate data for a sample species, the Fennec Fox (*Vulpes zerda*). a) Starting with range data from IUCN and climate data from ERA5 or an ESM, temperature and precipitation information is computed and the species range is annotated. b) The local climate limits for each grid-cell are computed over the species range. c) These are then aggregated into species niche limits over the range, representing 3 aspects - maximum/minimum of temperature/precipitation, variability in temperature/precipitation, and extreme events. The full list of species climate limits and calculation methodologies are detailed in Table 1.

### 2.1 Species range data

We sourced species range data from expert range maps (ERMs) as shapefiles from the IUCN Red List (IUCN, 2025) for all terrestrial and freshwater animal groups except birds. We sourced ERMs for all bird species from BirdLife International (Birdlife, 2024). Our dataset thus includes the resident or breeding ranges of all native mammals, amphibians, reptiles, and freshwater groups of fish and invertebrates. These ERMs represent curated range maps for each species as inferred by experts in different taxa, and thus represent an accurate assessment of a species’ geographic range.

Note that for many species, IUCN also provides resolutions of the range at the level of individual populations living in different areas. Given genetic differences and variation in environmental conditions across a species’ range, different populations may have different climate limits. We estimated climate limits at the level of each population separately, and aggregated the population-level limits to the species-level in our final database.

We assigned a realm to each species (i.e. terrestrial, freshwater, or terrestrial-freshwater) based on classifications provided by IUCN in the species range shapefiles. For species annotated as both terrestrial and freshwater (i.e. all amphibians, many reptiles, and other species whose life cycle or diet depends on water), we retained this classification. We annotated the range of each population using a 24km x 24km equal-area grid in EPSG:6933 (Brodzik et al., 2012), producing 875,520 grid-cells over the Earth’s surface (Figure 1a), and identifying the grid-cells occupied by a species’ range shapefile polygon.

Since we use static species ranges from modern-day records (ranging from 2005-2024) coupled with a time-series of climate over the last 80 years, we carry out an additional calculation to quantify the scale of the biasing effect that could come from species range shifts over 80 years given estimated rates from the BioShifts database (Lenoir et al., 2020). This analysis can be found in Supplementary Material S5; from this, we find that our estimated species climate limits are largely robust to range shifts for most species given the rate of shift and the scale of spatial autocorrelation in climate variables.

For each species in our dataset, we also collate information of taxonomic and conservation relevance, namely the kingdom, phylum, class, order, family, and IUCN Red List status. Further information on specific filtering steps is available in the Supplementary Material Section S1.1.

### 2.2 Historical climate data

Data on historical climate conditions of daily precipitation (pr), maximum daily temperature (tasmax), and minimum daily temperature (tasmin) were sourced from the ERA5 reanalysis project (Hersbach et al., 2023) representing interpolated observational climate data at a 0.25 degree resolution, and five bias-corrected Earth System Models (ESMs) from ISIMIP3b (Inter-Sectoral Impact Model Intercomparison Project) climate data records: EC-EARTH3, IPSL-CM6A-LR, GFDL-ESM4, MIROC6, and MPI-ESM1-2-HR (Lange, 2019) (Fig. 1a) at a 0.5 degree resolution. The sourcing and filtering of this data are described in the Supplementary Material Section S1.2. We use time periods of 1940-2020 for the ERA5 model, and 1941-2014 for the ISIMIP historical ESM projections, depending on the time periods available for each record. Among all 6 climate records, ERA5 is likely the most accurate approximation of past global real-world conditions available, as it is based on the reanalysis of real-world empirical observations. We used daily maximum/minimum temperature but aggregated precipitation data at the monthly scale using the `monsum` function from the cdo program (Schulzweida, 2023) since daily precipitation is usually 0 in most locations on most days. We also aggregated precipitation data into annual precipitation using the `yearsum` function from cdo. We reprojected the climate rasters into the same 24km x 24km equal-area grid-cells (in EPSG:6933) used for the species range shapefiles by bilinear interpolation (`remapbil` function from cdo).

### 2.3 Local climate limits (24×24 km grid-cell level)

We defined a local climate limit as a parameter representing the most extreme values seen in a single grid-cell computed from historical climate data for each of the 875,520 24km x 24km grid-cells (Fig. 1b). We estimated a total of 44 local climate limits split into 3 aspects: a) minima/maxima of historical daily temperature and of monthly/annual precipitation based on percentiles (26 climate limits), b) maxima of annual variability of temperature and of precipitation (4 climate limits), and c) maxima in the properties of extreme events (14 climate limits).

Specifically, temperature and precipitation maxima/minima represent the highest/lowest conditions experienced in a grid-cell defined by a given percentile. We selected percentiles of 95, 99, 99.9 and 100 (and correspondingly 5, 1, 0.1 and 0), to cover a range of threshold conditions as well as to buffer against potential outliers in the global climate records. Additionally, we also computed seasonal maximum/minimum temperature based on the warmest/coldest window of the year - we did this by constructing a climatological curve indicating the value of a specified percentile of temperature values from all years in a time-window of 10 days around a specified day (Hobday et al., 2016), then taking the maximum/minimum as the temperature in the warmest/coldest window. Annual variability in temperature and precipitation in a grid-cell is quantified by the coefficient of variation and the standard deviation. Extreme events represent sustained periods of extreme supra-threshold conditions in temperature and precipitation, in the form of heatwaves, cold-spells, and droughts.

We defined two types of extreme temperature events (Fig. S1a and S1b). In the first type, termed as heatwaves (and cold-spells), we use the definition of heatwaves as periods of 5 days or more when the temperature exceeds local limits from the climatology curve for that time of year (Hobday et al., 2016). In the second type, termed as maximum heatwaves (and minimum cold-spells), we defined them as periods of 5 days or more when the temperature breaches the maximum (or minimum) value across the entire climatology, regardless of any specific period of the year. This second type of extreme event represents a sustained period where conditions are more extreme than the warmest/coldest period of the year (more details in Suppl. S2.3).

We considered only droughts as extreme precipitation events, but not floods. Properties of droughts were calculated using the SPEI (Standardised Precipitation Evapotranspiration Index) (Vicente-Serrano et al., 2010), a multi-scale drought index incorporating temperature and geographical information with precipitation to measure expected periods of extremely low precipitation, with a timescale of 3 months, relevant to definitions of ecological drought (Crausbay et al., 2017). A drought starts when SPEI values fall below -1 for at least two consecutive months and ends when the index returns to a positive number (Medri et al., 2020) (Fig. S1c). Grid-cells with low vegetative coverage and strong precipitation seasonality, where the SPEI occasionally produced infinite values, were excluded from the analysis- as part of this, 121 species whose ranges were entirely comprised of such grid-cells were not included in the calculation of drought-related climate limits.

In Section S2 of the Supplementary Material, we describe in detail the computation of all local climate limits, and their potential ecological relevance.

### 2.4 Species climate limits (geographical-range level)

Species climate limits are defined as the most acute conditions experienced by the species over its geographic range; they represent the boundaries of the species’ realized niche in each of the 44 climate limits we considered. Species climate limits are computed by aggregating the values of local climate limits in each grid-cell of the species’ entire range (Fig. 1c). Specifically, species climate limits were estimated by taking the maximum/minimum of each local climate limit (Section 2.3) over the species’ complete geographical range. For example, the species climate limit T*_max_*_99_ is computed for a focal species by first finding the local climate limits of the 99th percentile of maximum daily temperature for all grid-cells the species is found in. Then, the species climate limit is the maximum of all these local climate limit values. In this way, we computed 44 species climate limits as realized limits over the window of 1940-2020 from all of the 44 local climate limits over the species’ entire range. Table 1 summarises the estimated 44 species climate limits produced for the ClimLimits database. Note that many species climate limits are correlated, representing similar physical quantities or a single limit computed at different cut-off percentiles for the local climate limits to account for different thresholds. Correlations between climate limits are presented in Suppl. S3.1. Additionally, we provide an option to compute regional-scale species climate limits for a specified geographic area. This is described in Suppl. S3.2.

A majority of species climate limits are calculated from commands in the cdo program, version 2.0.4 (Schulzweida, 2023) version 2.0.4. Seasonal climatology-based metrics and extreme events are computed using the ‘heatwaveR’ package (version 0.5.4) in R (Schlegel and Smit, 2018). All analyses were carried out in R version 4.3.3 (Team et al., 2024). Minor optimizations of the R scripts to improve runtime and performance were done using ChatGPT-4 (OpenAI, 2025).

## 3 Results

### 3.1 Database Records

The ClimLimits database consists of realized species climate limits for 54,255 species (80,016 populations) of terrestrial and freshwater taxa groups spanning all terrestrial biomes (Fig. 2a), comprising 24,731 terrestrial, 18,182 freshwater, and 11,342 terrestrial/freshwater species. We estimated the richness at the grid-cell level as the number of species whose ranges are in that grid-cell. Fig. 2b depicts the coverage of the different taxonomic groups in the ClimLimits database, with a majority of them (23.9%, 12,947 species) being freshwater fish. Reptiles (17.3%, 9,360 species) and birds (17%, 9,207 species) make up the two other major groups. Other freshwater groups comprise 5.2% of the database, and include crayfish, crabs, and shrimp. A large majority of species (40,018; 73.8%) have only one population annotated by the IUCN shapefiles over their range (Fig. 2c). Only 0.3% of species have more than 10 populations - these are typically cases of insular species whose island populations are annotated independently.

**Figure 2:**
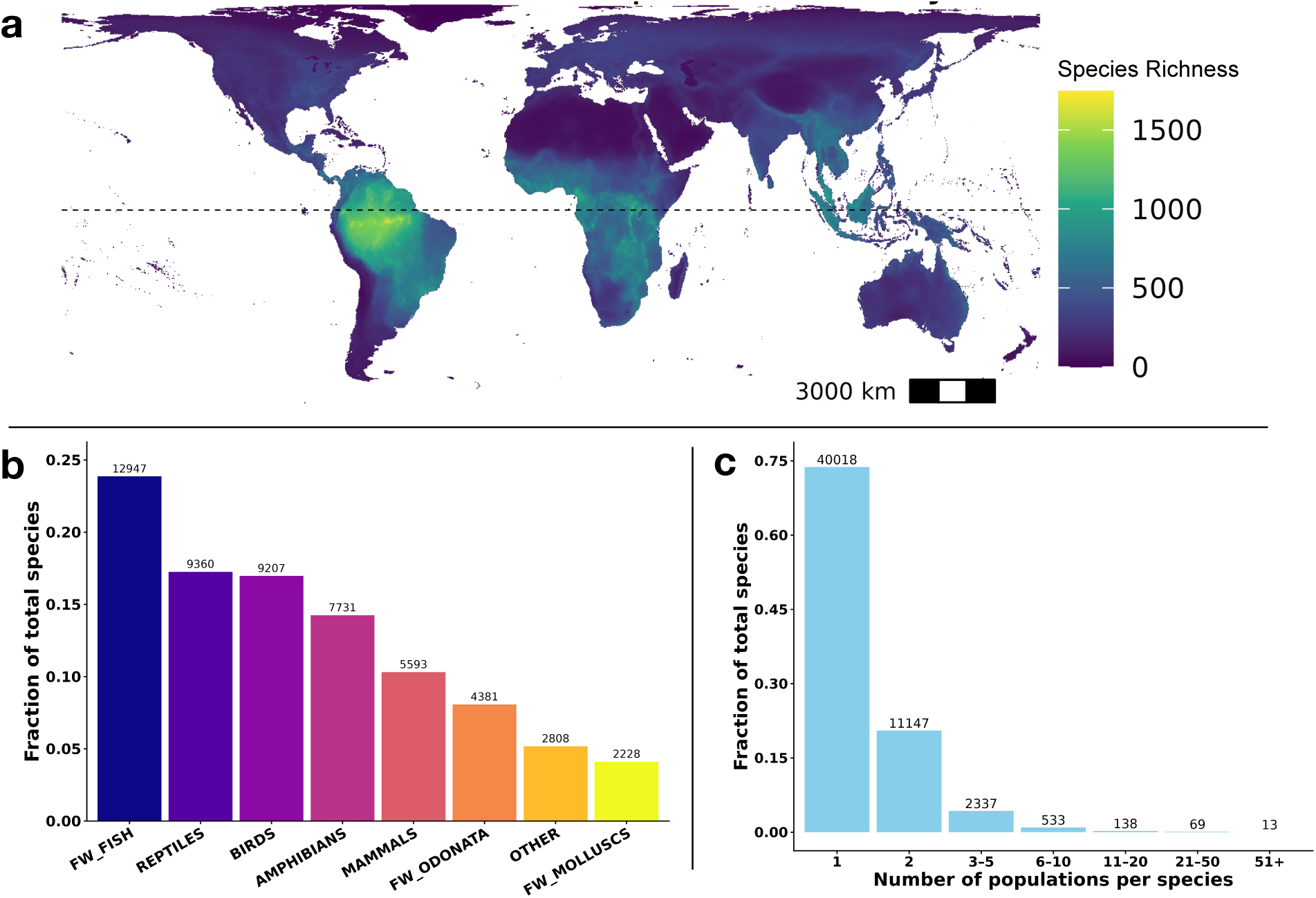
Summary of ClimLimits database coverage. (a) Global distribution of species richness in the ClimLimits database. Hotspots of diversity are visible in northwestern Amazonia, sub-Saharan Africa, and South East Asia, concordant with real-world patterns of biodiversity. (b) Fraction of total species per group of taxa in the ClimLimits database. Numbers above the bars correspond to the number of species per taxonomic group. Freshwater fish (FW_fish) are best represented, while reptiles, birds, amphibians, mammals, and freshwater odonates also contribute a large amount of diversity. The “other” category includes all other freshwater taxa, including freshwater crayfish, crabs, and shrimp. (c) Fraction of species in the ClimLimits database with one or more populations, with associated number of species above the bars. The IUCN shapefiles sometimes include multiple separate populations per species - we observe that 73.7% of species have only one population, with the remaining mostly have less than ten subpopulations.

The main database is presented in the form of seven .csv files. Six files contain estimates of the 44 species climate limits for 54,255 species for each of the six historical climate records-the ERA5 reanalysis, and five ISIMIP3b bias-corrected ESMs - EC-EARTH3, IPSL-CM6A-LR, GFDL-ESM4, MIROC6, and MPI-ESM1-2-HR. Each of these files also includes, for each species: i) the type of habitat of the species (terrestrial, freshwater or both), ii) the source shapefile from the IUCN/BirdLife international records, iii) the range size (expressed as the number of grid-cells the species is found in), iv) the latitudinal extent (the latitudinal distance over which the species’ range spans), v) the year that the species’ range polygon was annotated in the shapefiles (ranging from 2005 to 2024), vi) taxonomic information on the species, including its kingdom, phylum, class, order, and family, and vii) the IUCN red list status of the species (where available). The seventh file is the `variable_metadata.csv` file that explains the variable names in the files representing the species’ climate limits. In addition to these seven files, the database contains two auxiliary files - one file contains information on the list of grid-cells each population is found in (`population_level_ranges_full.csv`), and another file maps each grid-cell index number to a latitude and longitude value (`boundaries_grid_cells_full.csv`). These are provided for reference and validation, as well as for visualizing species ranges on a global map (see Section S4).

### 3.2 Case Study: Modeling Species Exposure Across Climate Limits in an SSP Scenario

To illustrate a potential use case of this database, we use the realized species climate limits from the ClimLimits database as a baseline to model future species exposure across multiple climate limits in a sample climate warming scenario over the coming century. This method is in line with recent studies assessing the scale of thermal exposure for species worldwide based on future climate scenarios (Trisos et al., 2020; Pigot et al., 2023), but uses a larger pool of climate limits beyond maximum temperature, and is computed at a higher spatiotemporal resolution (daily-scale across 24km x 24km grid-cells) to reflect more accurately the most extreme conditions that species have historically experienced. Crucially, the multivariate nature of ClimLimits lets us compare the extent of exposure across climate limits that will be breached the most for a single species or across global ecosystems, identifying which factors are likely to be the largest drivers of future risk.

To do this, we source sample bias-corrected future projections of daily maximum/minimum temperature and monthly/yearly precipitation from the SSP5-8.5 scenario for the MPIESM model from the ISIMIP3b project (Lange, 2019). The SSPs (Shared Socioeconomic Pathways) represent potential future climate trajectories based on socioeconomic choices made by the human population over the coming century (2021-2100); SSP5-8.5 represents the worst of these, with a heavy reliance on fossil fuels leading to an average warming of roughly 4K above pre-industrial conditions. We select the last 20 years of future projection data (2081-2100), and compute the local climate limits for each grid-cell using the same procedure as that used for ClimLimits (described in Section 2.3). Then, we select 16 climate limits from the pool of 44: T*_max_*_99_, T*_min_*_1_, Pr*_max_*_99_, Pr*_min_*_1_, YPr*_max_*_99_, YPr*_min_*_1_, CV*_temp_*, CV*_pr_*, HW*_freq_*, CS*_freq_*, maxHW*_freq_*, maxHW*_maxdur_*, maxHW*_maxint_*, DR*_f req_*, DR*_maxint_*, and DR*_maxdur_* (see Table 1 for definitions of each of these). For each grid-cell in a focal species’ range, we define exposure if the value of the local climate limit in the future exceeds the realized historical species climate limit. For example, if the future local climate limit for T*_max_*_99_ (99th percentile of maximum temperature) in a cell is 323K, while the historical species climate limit for T*_max_*_99_ was 321K, we consider this species to be exposed in this cell. This way, for each species and each of the 16 climate limits, we can find the fraction of the species’ range exposed in this scenario of future climate warming. Though this is a simplistic binarized assessment of exposure without accounting for the extent of the exposure and the pre-existing suitability of the grid-cell for the species, this allows us to get preliminary assessments of exposure risk and the comparative effect across variables.

We identify exposure percentages per climate limit for all species with range size greater than 10 grid-cells (n = 43,230). Figure 3 depicts the percentage of each species’ range exposed in each climate limit, divided into 3 broad categories: a) full exposure, meaning that all grid-cells in the species range have future local climate limits exceeding the historical species climate limit; b) partial exposure, meaning that some, but not all grid-cells are exposed; and c) no exposure, indicating that no grid-cells have their future local climate limits exceeding the species climate limit. We find that in the MPIESM SSP5-8.5 scenario, exposure is pervasive across most climate limits, with partial and full exposure for a large number of species in many climate limits. However, we note that properties pertaining to heatwaves (especially heatwave frequency and maximum heatwave frequency) have a large fraction of species exposed over their entire range, corresponding to 40,515 and 32,984 species for heatwave frequency and maximum heatwave frequency). This indicates that in this scenario, a large majority of species are experiencing more frequent heatwaves over their entire geographic range than the highest frequency they had seen in the past, illuminating heatwave frequency as a major driver of future biodiversity vulnerability. We highlight this as a unique multivariate assessment of comparative risk from different aspects of future climate change that can be done by using the ClimLimits database.

**Figure 3:**
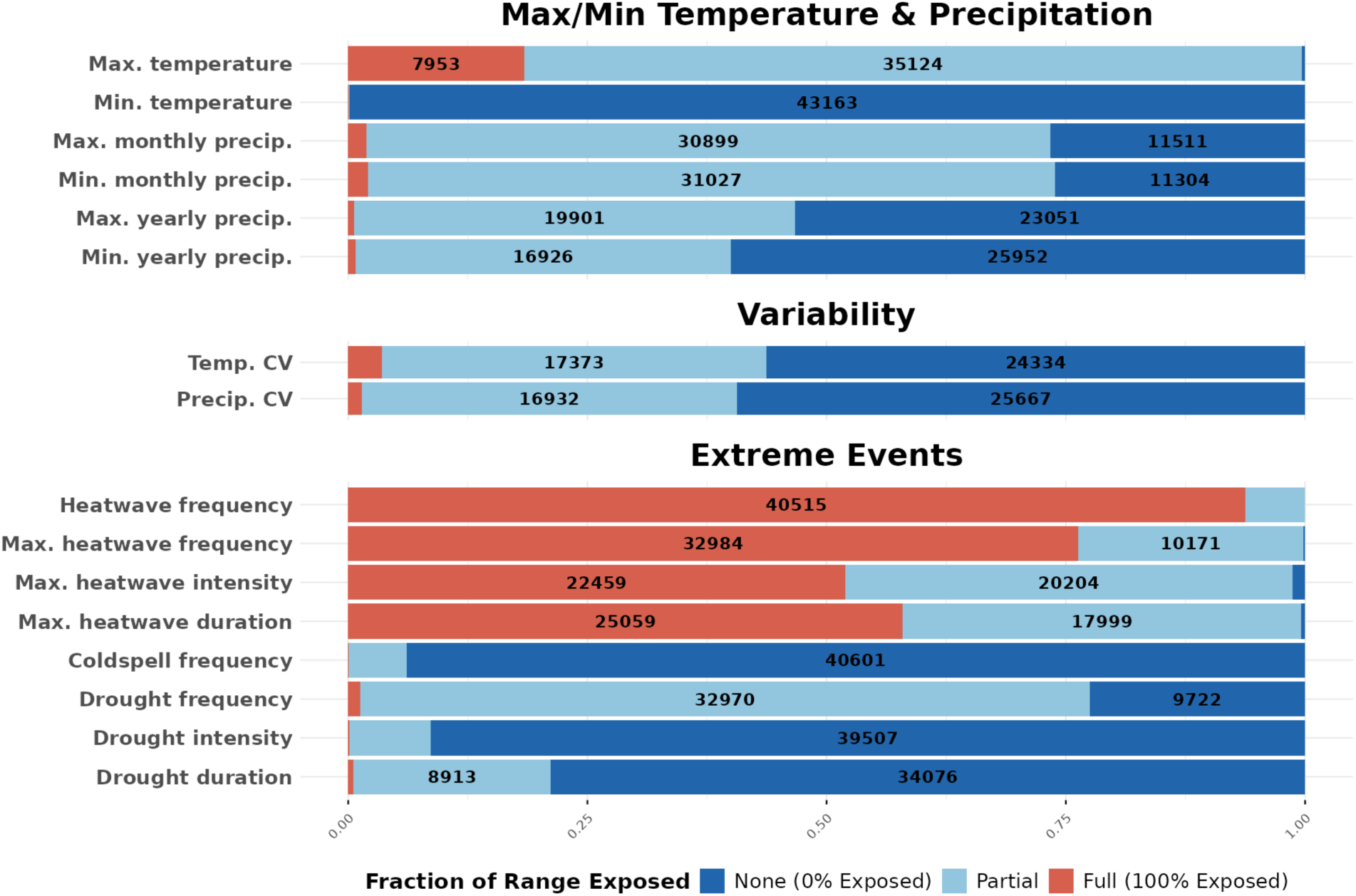
Case study: fraction of species with geographical ranges not, partially or fully exposed to 16 climate limits under the MPIESM SSP5-8.5 projected climate scenario in the time period 2081-2100. Exposure is defined as fraction of grid-cells with local climate limits exceeding the species’ climate limit. The percentages of species geographical ranges exposed to a given climate limit are grouped into three categories: not exposed (in light blue), partially exposed (in dark blue), and fully exposed (in red). The numbers in the colored boxes correspond to the number of species (out of 43,230) not exposed, partially exposed or fully exposed to each climate limit.

Additionally, this exposure risk can be quantified in other aspects as well - this same technique can be extended to identify geographic regions of highest exposure. This can be done by selecting the assemblage of species found in a single grid-cell and assessing how many of them are exposed in that particular grid-cell for a given climate limit, generating a spatial map of exposure extent. Figure 4 shows the number of species exposed per grid-cell on average across all 16 climate limits selected. This reveals that the number of species exposed is the highest around the Amazon region and parts of Southeast Asia, suggesting that these regions are likely to pose the most stressful conditions on average for species in the future and therefore may be targets for climate change mitigation and conservation measures.

**Figure 4:**
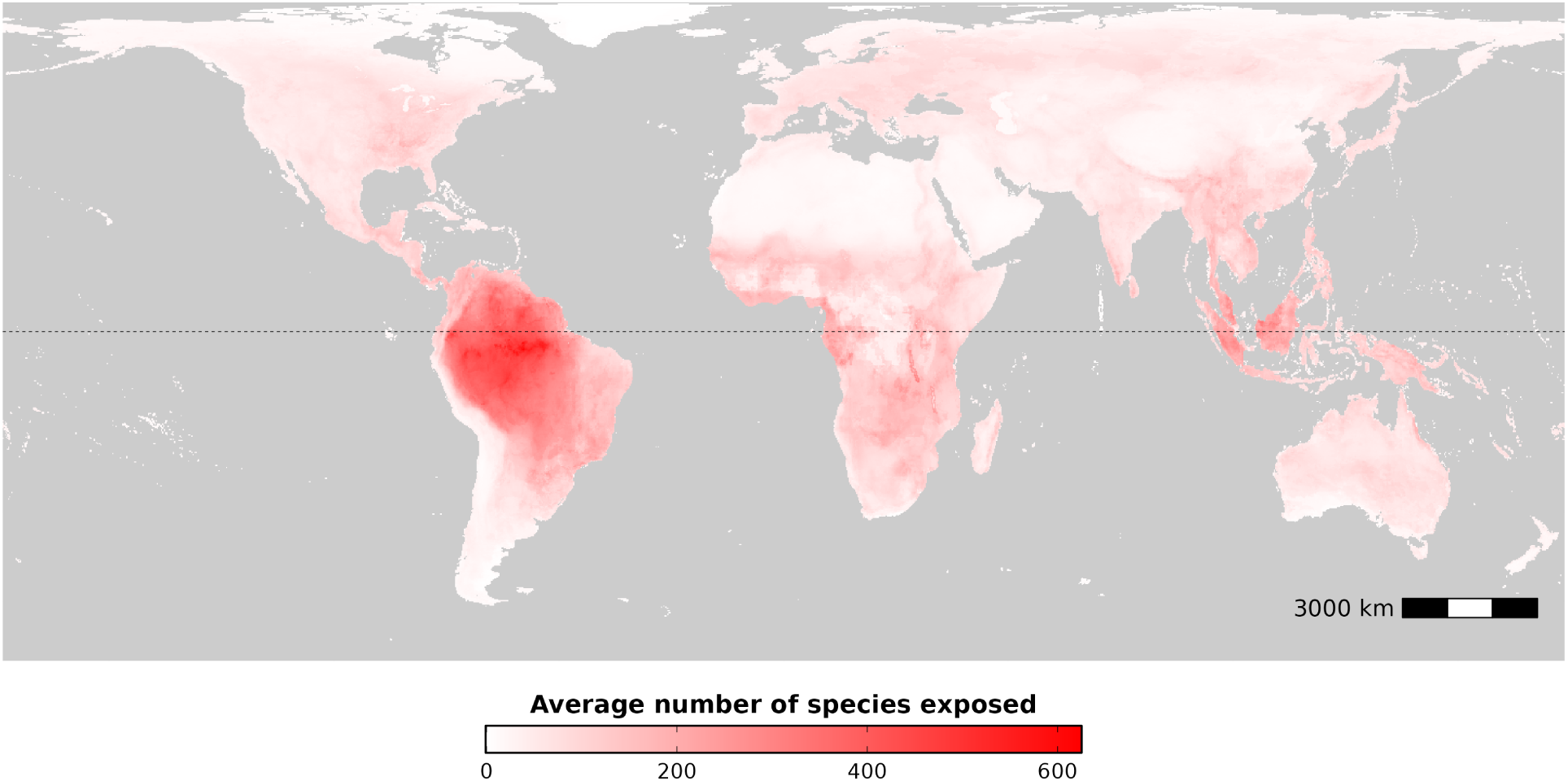
Case study: average number of species exposed per grid-cell in 2081-2100 to 16 climate limits per grid-cell under the MPIESM SSP5-8.5 projected climate scenario. Exposure is computed here at the level of species assemblage in a given grid-cell, then averaged over all 16 climate limits to identify spatial patterns of risk.

## 4 Discussion

### 4.1 Conservation Implications

The ClimLimits database is designed as a resource for macroecological assessments of patterns of species climate limits and modeling climate change exposure risk at a high spatiotemporal resolution, at both regional and global scales. In particular, it accounts for multiple aspects of climate, including variability and daily-scale climate extremes, aspects frequently overlooked in determining climate impacts on biodiversity (Ma et al., 2015). We envision several uses of the ClimLimits database, ranging from climate change sensitivity and exposure assessments, to macroecological syntheses and identification of regions and species at maximum risk of high exposure and conservation priority. As indicated in the above case study (Section 3.2), ClimLimits can be used to compute exposure patterns and compare drivers of exposure in any given climate scenario, and for a particular set of species or geographic regions.

ClimLimits builds on approaches taken by recent papers assessing exposure by modeling exceedance of species’ realized historical temperature limits (Trisos et al., 2020; Murali et al., 2023; Pigot et al., 2023; Meyer et al., 2024; Watson and Kerr, 2025). Compared with previous studies, ClimLimits allows for assessments beyond previously-used univariate (or bivariate) approaches and coarse spatiotemporal resolution (such as modeling climate limits in 100km × 100km grid-cells using monthly or annual average temperature). In that sense, ClimLimits provides a standardized, reproducible and high-resolution record of realized climate limits across historical climate records, temperature/precipitation variables, geographical regions, and taxa.

### 4.2 Comparison with other climate limits

The ClimLimits database also complements other climate-limit databases, such as GlobTherm (Bennett et al., 2018) or ThermoFresh (Bayat et al., 2025). While these databases include experimentally-measured estimates of species’ fundamental niche limits, ClimLimits provides realized climate niche limits inferred from species ranges and associated historical climate conditions. The use of species range and climate data by ClimLimits permits coverage of a much larger number of species and taxa (n = 54,255) compared to Thermofresh (n = 931) or Globtherm (n = 2,133). When comparing limits in ClimLimits to the ones from GlobTherm for cases where data is available, we find that there is a fair degree of correlation- the Pearson correlation coefficient between thermal maxima is 0.32 (p < 2.2e-16, n = 673 species), indicating that these agree reasonably well. However, it is important to note the caveat that these represent intrinsically different quantities, with many possible factors influencing one to be either higher or lower than the other. For example, biotic interactions and dispersal limitations would lead to the realized limit from ClimLimits being lower than the experimentally-measured fundamental limit, while unaccounted-for fine-scale microclimatic refugia in climate records may cause the realized limit to be higher (Chevalier et al., 2024).

At the same time, the multivariate nature of ClimLimits adds to the recent database of seasonally-varying realized limits in maximum/minimum aridity and temperature for terrestrial vertebrates (Watson and Kerr, 2025). Our database computes a larger pool of climate limits including properties of variability and extreme events based on daily-scale extremes, covering a longer historical timescale, and expanding the species pool covered (from 33,941 to 54,255) to include freshwater taxa.

### 4.3 Potential Biases

Potential biases in the estimates of the species climate limits of our database may arise due to: (a) taxonomic and spatial bias arising from differing range quality annotations, especially for understudied species in the Global South; (b) microclimatic and topographic variation in temperature modifying species climate limits beyond what can be captured at a 24km x 24km scale resolution (as cautioned in (Colwell, 2021; Chevalier et al., 2024)); (c) potential systematic errors arising from the use of global climate records, especially in the tropics (see (Farneti et al., 2022)); (d) the assumption that there is no local adaptation that could make certain populations less tolerant to the climate limits measured as the global maxima/minima over the range; and (e) the assumption of species’ equilibrium with their climate (Araújo et al., 2005) and that their ranges are static over the entire historical period, not moving towards their optimal range (but see Supplementary Material S5). Additionally, biases in our estimates of species climate limits could be especially prominent for species which have undergone recent large-scale habitat loss due to anthropogenic activity and land-use change (for example, the grey wolf (*Canis lupus*))- these species may have a significantly larger historical range than what is recorded by recent IUCN shapefiles and therefore will likely have wider climate limits than our database estimates. However, over a species’ current range, our species climate limits provide a conservative estimate of the species’ realized limits, given that we select the global maximum over all cells, despite there being local populations that may have comparatively lower climate limits.

### 4.4 Future directions and application

The reproducible structure of ClimLimits allows our method to be easily updated with any historical climate model output, and readily adapted to work with any record of species ranges such as GBIF (GBIF, 2025), outputs from Species Distribution Models (SDMs) at global or regional scales (Jetz et al., 2012; Adde et al., 2025), or databases for specific taxa such as the Phylacine database or eBird (for mammals and birds, respectively) (Sullivan et al., 2009; Faurby et al., 2018). Producing species climate limits from alternative climate and range map sources may reduce some of the aforementioned biases, and could account for different model uncertainties, species sampling biases, and range annotation accuracies.

Therefore, the ClimLimits database presents a standard set of values alongside a repro-ducible general framework of climate limit estimation to synthesize multivariate realized species climate limits in an accessible and high-resolution format, facilitating detailed analyses of macroecological patterns and species vulnerability assessments in the present and future.

## Author contributions

K.S.G., V.D., and C.J. conceptualized the study. V.D. and C.J provided resources and supervision, acquired funding and handled project administration. Data curation, formal analysis, software development, and validation were carried out by K.S.G. Investigation and visualization were done by K.S.G., V.D., and C.J. K.S.G. wrote the original draft of the manuscript, V.D. and C.J. provided revisions and edits to the manuscript.

## Acknowledgments

The authors would like to thank Iago Bonnici, Miguel Berdugo, Philipp Hess, Santiago Beguería, and Sebastian Bathiany for assistance with processing climate data, and the Université de Montpellier cantine (except on the days they have dal) for providing the energy to write this.

## Data and code availability statement

.csv files constituting the database, as well as the R scripts used to produce the files, are available on Zenodo at the following link: https://doi.org/10.5281/zenodo.21397317 (currently anonymized for peer-review). Intermediate datasets that may be required for specific further analyses but may not be possible to reproduce due to computing limitations may be available on request to the corresponding author.

## Supporting information

Additional supporting information may be found online.

## References

Adde, A., Rey, P.-L., Külling, N., Chauvier-Mendes, Y., Fopp, F., Popp, M. R., Broennimann, O., Petitpierre, B., Strebel, N., Gross, A., Stofer, S., Lehmann, A., Zimmermann, N. E., Pellissier, L., Guisan, A., and Altermatt, F. (2025). SDMapCH: A Comprehensive database of *>*7,500 modelled species habitat suitability maps for Switzerland. Scientific Data, 12(1):1752.

Araújo, M. B., Pearson, R. G., and Rahbek, C. (2005). Equilibrium of species’ distributions with climate. Ecography, 28(5):693–695.

Bathiany, S., Dakos, V., Scheffer, M., and Lenton, T. M. (2018). Climate models predict increasing temperature variability in poor countries. Sci. Adv., 4(5):eaar5809.

Bayat, H. S., He, F., Medina Madariaga, G., Escobar-Sierra, C., Prati, S., Peters, K., Jupke, J. F., Spaak, J. W., Manfrin, A., Juvigny-Khenafou, N. P. D., Chen, X., and Schäfer, R. B. (2025). Global thermal tolerance compilation for freshwater invertebrates and fish. Scientific Data, 12(1):1488.

Bennett, J. M., Calosi, P., Clusella-Trullas, S., Martínez, B., Sunday, J., Algar, A. C., Araújo, M. B., Hawkins, B. A., Keith, S., Kühn, I., Rahbek, C., Rodríguez, L., Singer, A., Villalobos, F., Ángel Olalla-Tárraga, M., and Morales-Castilla, I. (2018). GlobTherm, a global database on thermal tolerances for aquatic and terrestrial organisms. Scientific Data, 5(1):180022.

Birdlife, I. (2024). Birdlife | partnership for nature and people.

Brodzik, M. J., Billingsley, B., Haran, T., Raup, B., and Savoie, M. H. (2012). EASE-Grid 2.0: Incremental but Significant Improvements for Earth-Gridded Data Sets. ISPRS International Journal of Geo-Information, 1(1):32–45.

Chevalier, M., Pignard, V., Broennimann, O., and Guisan, A. (2024). A cautionary message on combining physiological thermal limits with macroclimatic data to predict species distribution. Ecosphere, 15(7).

Colwell, R. K. (2021). Spatial scale and the synchrony of ecological disruption. Nature, 599(7886):E8–E10.

Crausbay, S. D., Ramirez, A. R., Carter, S. L., Cross, M. S., Hall, K. R., Bathke, D. J., Betancourt, J. L., Colt, S., Cravens, A. E., Dalton, M. S., Dunham, J. B., Hay, L. E., Hayes, M. J., McEvoy, J., McNutt, C. A., Moritz, M. A., Nislow, K. H., Raheem, N., and Sanford, T. (2017). Defining ecological drought for the twenty-first century. Bull. Am. Meteorol. Soc., 98(12):2543–2550.

De Luca, P. and Donat, M. G. (2023). Projected changes in hot, dry, and compound hot-dry extremes over global land regions. Geophys. Res. Lett., 50(13).

Farneti, R., Stiz, A., and Ssebandeke, J. B. (2022). Improvements and persistent biases in the southeast tropical atlantic in cmip models. npj Climate and Atmospheric Science, 5(1):42.

Faurby, S., Davis, M., Pedersen, R. Ø., Schowanek, S. D., Antonelli, A., and Svenning, J.-C. (2018). Phylacine 1.2: the phylogenetic atlas of mammal macroecology. Ecology, 99(11):2626.

Freeman, B. G., Scholer, M. N., Ruiz-Gutierrez, V., and Fitzpatrick, J. W. (2018). Climate change causes upslope shifts and mountaintop extirpations in a tropical bird community. Proceedings of the National Academy of Sciences, 115(47):11982–11987.

Gao, Y., Wu, Y., Guo, X., Kou, W., Zhang, S., Leung, L. R., Chen, X., Lu, J., Diffenbaugh, N. S., Horton, D. E., Yao, X., Gao, H., and Wu, L. (2023). More frequent and persistent heatwaves due to increased temperature skewness projected by a high-resolution earth system model. Geophys. Res. Lett., 50(18).

Garcia, R. A., Cabeza, M., Rahbek, C., and Araújo, M. B. (2014). Multiple Dimensions of Climate Change and Their Implications for Biodiversity. Science, 344(6183):1247579.

Garnier, E., Pelletier, D., Roche, P., Rougerie, R., and Pavoine, S. (2025). Defining biodiversity data. Trends in Ecology & Evolution, 40(8):731–735.

GBIF (2025). GBIF — gbif.org. https://www.gbif.org. [Accessed 11-12-2025].

Gunderson, A. R. and Stillman, J. H. (2015). Plasticity in thermal tolerance has limited potential to buffer ectotherms from global warming. Proceedings of the Royal Society B: Biological Sciences, 282(1808):20150401.

Halpern, B. S., Frazier, M., Potapenko, J., Casey, K. S., Koenig, K., Longo, C., Lowndes, J. S., Rockwood, R. C., Selig, E. R., Selkoe, K. A., and Walbridge, S. (2015). Spatial and temporal changes in cumulative human impacts on the world’s ocean. Nature Communications, 6(1):7615.

Hersbach, H., Bell, B., Berrisford, P., Biavati, G., Horányi, A., Muñoz Sabater, J., Nicolas, J., Peubey, C., Radu, R., Rozum, I., Schepers, D., Simmons, A., Soci, C., Dee, D., and Thépaut, J.-N. (2023). Era5 hourly data on single levels from 1940 to present.

Hobday, A. J., Alexander, L. V., Perkins, S. E., Smale, D. A., Straub, S. C., Oliver, E. C., Benthuysen, J. A., Burrows, M. T., Donat, M. G., Feng, M., Holbrook, N. J., Moore, P. J., Scannell, H. A., Sen Gupta, A., and Wernberg, T. (2016). A hierarchical approach to defining marine heatwaves. Progress in Oceanography, 141:227–238.

IUCN (2025). The iucn red list of threatened species.

Jetz, W., McPherson, J. M., and Guralnick, R. P. (2012). Integrating biodiversity distribution knowledge: toward a global map of life. Trends in Ecology & Evolution, 27(3):151–159.

Karger, D. N., Saladin, B., Wüest, R. O., Graham, C. H., Zurell, D., Mo, L., and Zimmermann, N. E. (2023). Interannual climate variability improves niche estimates for ectothermic but not endothermic species. Sci. Rep., 13(1):12538.

Kefford, B. J., Nichols, S. J., and Duncan, R. P. (2023). The cumulative impacts of anthropogenic stressors vary markedly along environmental gradients. Global Change Biology, 29(3):590–602.

Kingsolver, J. G., Diamond, S. E., and Buckley, L. B. (2013). Heat stress and the fitness consequences of climate change for terrestrial ectotherms. Functional Ecology, 27(6):1415–1423.

Lange, S. (2019). Trend-preserving bias adjustment and statistical downscaling with ISIMIP3BASD (v1.0). Geoscientific Model Development, 12(7):3055–3070.

Lenoir, J., Bertrand, R., Comte, L., Bourgeaud, L., Hattab, T., Murienne, J., and Grenouillet, G. (2020). Species better track climate warming in the oceans than on land. Nature ecology & evolution, 4(8):1044–1059.

Ma, G., Hoffmann, A. A., and Ma, C.-S. (2015). Daily temperature extremes play an important role in predicting thermal effects. Journal of Experimental Biology, 218(14):2289–2296.

Malhi, Y., Franklin, J., Seddon, N., Solan, M., Turner, M. G., Field, C. B., and Knowlton, N. (2020). Climate change and ecosystems: Threats, opportunities and solutions. Philosophical Transactions of the Royal Society B: Biological Sciences, 375(1794):20190104.

Medri, S., Crespi, A., Terzi, S., Cocuccioni, S., Zebisch, M., Berckmans, J., and Füssel, H.-M. (2020). Climate-related hazard indices for europe. Technical Report Technical Paper 1/2020, European Topic Centre on Climate Change Adaptation (ETC/CCA), Bologna, Italy. Prepared by EURAC Research, VITO, and the European Environment Agency (EEA).

Meyer, A. S., Pigot, A. L., Merow, C., Kaschner, K., Garilao, C., Kesner-Reyes, K., and Trisos, C. H. (2024). Temporal dynamics of climate change exposure and opportunities for global marine biodiversity. Nature Communications, 15(1).

Michener, W. K. and Jones, M. B. (2012). Ecoinformatics: Supporting ecology as a data-intensive science. Trends in Ecology & Evolution, 27(2):85–93.

Murali, G., Iwamura, T., Meiri, S., and Roll, U. (2023). Future temperature extremes threaten land vertebrates. Nature, 615(7952):461–467. OpenAI (2025). Chatgpt-4.

Parmesan, C. and Yohe, G. (2003). A globally coherent fingerprint of climate change impacts across natural systems. Nature, 421(6918):37–42.

Pearson, R. G., Thuiller, W., Araújo, M. B., Martinez-Meyer, E., Brotons, L., McClean, C., Miles, L., Segurado, P., Dawson, T. P., and Lees, D. C. (2006). Model-based uncertainty in species range prediction. Journal of Biogeography, 33(10):1704–1711.

Perkins, S. E. and Alexander, L. V. (2013). On the Measurement of Heat Waves. Journal of Climate, 26(13):4500–4517.

Pigot, A. L., Merow, C., Wilson, A., and Trisos, C. H. (2023). Abrupt expansion of climate change risks for species globally. Nature Ecology & Evolution, 7(7):1060–1071.

Schlegel, R. W. and Smit, A. J. (2018). heatwaveR: A central algorithm for the detection of heatwaves and cold-spells. Journal of Open Source Software, 3(27):821.

Schulzweida, U. (2023). Cdo user guide.

Sullivan, B. L., Wood, C. L., Iliff, M. J., Bonney, R. E., Fink, D., and Kelling, S. (2009). ebird: A citizen-based bird observation network in the biological sciences. Biological Conservation, 142(10):2282–2292.

Sunday, J. M., Bates, A. E., Kearney, M. R., Colwell, R. K., Dulvy, N. K., Longino, J. T., and Huey, R. B. (2014). Thermal-safety margins and the necessity of thermoregulatory behavior across latitude and elevation. Proceedings of the National Academy of Sciences, 111(15):5610–5615.

Team, R.-C. et al. (2024). R version 4.3. 3. r core team.

Thomas, C. D., Cameron, A., Green, R. E., Bakkenes, M., Beaumont, L. J., Collingham, Y. C., Erasmus, B. F. N., De Siqueira, M. F., Grainger, A., Hannah, L., Hughes, L., Huntley, B., Van Jaarsveld, A. S., Midgley, G. F., Miles, L., Ortega-Huerta, M. A., Townsend Peterson, A., Phillips, O. L., and Williams, S. E. (2004). Extinction risk from climate change. Nature, 427(6970):145–148.

Trisos, C. H., Merow, C., and Pigot, A. L. (2020). The projected timing of abrupt ecological disruption from climate change. Nature, 580(7804):496–501.

Troudet, J., Grandcolas, P., Blin, A., Vignes-Lebbe, R., and Legendre, F. (2017). Taxonomic bias in biodiversity data and societal preferences. Scientific Reports, 7(1):9132.

Trull, N., Böhm, M., and Carr, J. (2018). Patterns and biases of climate change threats in the IUCN Red List. Conservation Biology, 32(1):135–147.

Urban, M. C. (2015). Accelerating extinction risk from climate change. Science, 348(6234):571–573.

Valavi, R., Guillera-Arroita, G., Lahoz-Monfort, J. J., and Elith, J. (2022). Predictive performance of presence-only species distribution models: A benchmark study with reproducible code. Ecological Monographs, 92(1):e01486.

Vicente-Serrano, S. M., Beguería, S., and López-Moreno, J. I. (2010). A multiscalar drought index sensitive to global warming: The standardized precipitation evapotranspiration index. J. Clim., 23(7):1696–1718.

Watson, M. and Kerr, J. (2025). Global dataset for realized thermal and aridity niche limits for terrestrial vertebrates. Sci. Data, 12(1):1829.

Weaving, H., Terblanche, J. S., Pottier, P., and English, S. (2022). Meta-analysis reveals weak but pervasive plasticity in insect thermal limits. Nature Communications, 13(1):5292.

Wiens, J. J. (2016). Climate-Related Local Extinctions Are Already Widespread among Plant and Animal Species. PLOS Biology, 14(12):e2001104.

