## Supplementary Material for "ClimLimits: a global database of multivariate realized climate limits for animal species"

#### S1 Data sources and filtering

##### S1.1 Species Range Data

We sourced species range data as shapefiles from expert range maps (ERMs) from the IUCN Red List (IUCN, 2025) from the IUCN website for all terrestrial and freshwater animal groups, except birds. This includes mammals, amphibians, reptiles, and all freshwater groups. We also source marine species' shapefiles, but exclude them from the analysis. The reason is that sea-surface temperature (SST), which is the primary driver of marine ecosystems' temperature response (Roy et al., 1998) at a global scale is only available at a monthly averaged scale, which we consider to be too coarse to compute species climate limits. We also excluded plants (both freshwater and terrestrial) for two reasons - (a) expert range maps from the IUCN for plants are extremely limited, and were often produced many years ago, thus producing concerns of data resolution and reliability; and (b) the governing biotic timescales and processes for plants are likely significantly different from those of animals. Specifically, plants are likely to have much higher climatic lags and disequilibrium in the face of climate change on account of their limited dispersal ability (Svenning and Sandel, 2013; Alexander et al., 2018; Sandel et al., 2025) (but see (Boonman et al., 2025)). Furthermore, plants likely have a different set of climate limits that govern their limits of existence than animals - for example, variables such as frost-free days, soil water, growing degree days, and vapor pressure deficit have all been shown to have significant direct effects on plant function in terms of affecting their phenology, growth, and survival (Chuine, 2010; Grossiord et al., 2020; Wang et al., 2020; Fu et al., 2022), but may affect animals more indirectly. Indeed, many of these relevant climate limits for plants have already been quantified as extremes in the ETCCDI indices (Karl et al., 1999; Peterson and Coauthors, 2001).

For bird species, we retrieved ERMs from the BirdLife International website (Birdlife, 2024). ERMs provide an accurate assessment of a species' range based on curation of available data sources from surveys and models by experts on the taxa. For many species, IUCN provides shapefiles of ranges at the level of individual populations living in different areas. Environmental variation across the range of different populations suggests genetic differences between these, leading to variation in species climate limits across populations; while we computed species climate limits at the population level in the code, we aggregated them to the species level in the final database for easier interpretability. To account for possibly less accurate ranges that may not represent the current accuracy of sampling methods, we removed species whose ranges were annotated before 2005. Then, we filtered the shapefiles to include only those that represent permanent ranges - that is, cases where species are recorded as extant, either by native presence or reintroduction, and are present either year-round or in the breeding season.

Then, using the `st_within` and `st_covers` functions from the 'sf' package in R (version 1.0-19) (Pebesma, 2018; Pebesma and Bivand, 2023), we rasterized the shapefile of each population into grid-cells on an equal-area grid of resolution 24km x 24km, generated in EPSG:6933 (Brodzik et al., 2012), returning the set of grid cells that constitute the species' range. The `st_within` function was used to identify which grid-cells contain each of the points on the boundary of the shapefile, while the `st_covers` function identifies the grid-cells contained entirely within the shapefile. Though this method may potentially exclude parts of the range which do not contain a boundary point of the shapefile, nor completely lie within the shapefile, the extent of this error is minimal- we quantified this by checking the accuracy of annotation as the size of the grid-cell range compared to the area of the original shapefile, and got an accuracy of 98.7%. We provide these range annotations, as well as the latitudinal and longitudinal boundaries of each of the 875,520 grid cells as two auxiliary files. We highlight that these can be used for visualizing the species' range on the world map (see Section S4).

Then, based on the classification provided by the IUCN shapefiles, we assigned a realm to each species as some combination of terrestrial, freshwater, and marine (a single species may belong to more than one realm). Since marine species are outside the scope of our database, we excluded species classified exclusively as marine. For species annotated both as marine and as terrestrial or freshwater, we used the GSHHG land mask (Wessel and Smith, 1996) to identify which grid-cells are marine, and excluded species whose ranges include more than 50% marine cells. For species with separate populations that are classified as belonging to different realms based on their locations, we excised the species if any of its populations are marine; if there is a mix of terrestrial/freshwater/both populations, we classified them as 'terrestrial+freshwater'. For species annotated by the IUCN as both terrestrial and freshwater (i.e. all amphibians, many reptiles, and other species such as waterbirds whose life cycle or diet depends on water), we retained this classification. We provide this annotation for each species as one of the columns in the final species climate limit database. We also source taxonomic information on the kingdom, phylum, class, order, and family of the species, as well as the IUCN red list status of the species. For species in the IUCN shapefiles, this is provided directly and collated for all species considered. However, for birds from the BirdLife International dataset, this is not provided - thus, for these, we sourced this information where available from the AviList database (Rheindt et al., 2025). However, due to taxonomic changes, some species were not matched with records in AviList - for these, we standardized synonyms where available from the GBIF (GBIF, 2025) with the rgbif package (Chamberlain et al., 2026). Even after this, we were left with 738 species (805 populations) which lacked a taxonomic match in AviList (typically due to more recent taxonomic changes not yet reflected in the rgbif call). For these species, we source the taxonomic information from the GBIF and do not report the IUCN red list status. This produced a final list of 54,255 species, representing 80,016 populations.

#### S1.2 Historical Climate Data

We sourced daily-scale data for tasmax (maximum air-surface temperature), tasmin (minimum air-surface temperature), and pr (total precipitation) from the ERA5 reanalysis (Hersbach et al., 2023) (accessed from the Copernicus server for ERA5) at a resolution of 0.25 degrees, and five bias-corrected ISIMIP3b (Inter-Sectoral Impact Model Intercomparison Project) models (Lange, 2019) (accessed from the ISIMIP website)- namely, EC-EARTH3, IPSL-CM6A-LR, GFDL-ESM4, MIROC6, and MPI-ESM1-2-HR, at a resolution of 0.5 degrees. We selected these five models on the basis of their availability of daily-scale bias-corrected data of the three variables of our interest. For ERA5 data, climate rasters for the variables '2m temperature' and 'total precipitation' were downloaded from the ERA5 server at an hourly scale for all days and all hours from the years 1940-2020 by making a call to the Copernicus API. Then, using the `daymax` and `daymin` functions from the cdo program (Schulzweida, 2023), we aggregated the temperature data into daily-scale tasmax and tasmin respectively. Similarly, we applied the `daysum` function from cdo to compute daily-scale precipitation. For ISIMIP models, we sourced the daily-scale tasmax, tasmin and precipitation data from the period of 1941-2014 as .txt files from the ISIMIP site providing the URL of the raster for each model. We computed all local climate limits from the three climate rasters of daily maximum temperature, daily minimum temperature, and daily total precipitation.

We opted to generate species climate limits from multiple climate records for three reasons. First, given the different modeling and simulation frameworks used to produce these models (Hawkins and Sutton, 2011; Zelinka et al., 2020), studies based on ESMs outputs frequently use an ensemble of models, producing a range of outputs (Tebaldi and Knutti, 2007; Eyring et al., 2019), which are either presented as a range of values or a single consensus value, as in, for example, (Garcia et al., 2012). This also supports a recent review highlighting the need for model intercomparison studies for biodiversity (Zurell et al., 2026). Second, the presence of systematic biases in the models (Flato et al., 2014) prevents easy comparison between them. ISIMIP3b models, however, are bias-corrected (Lange, 2019), while the ERA5 reanalysis record (Hersbach et al., 2023) is fitted to observational data, improving its accuracy. Thirdly, using ESMs from ISIMIP3b allows for having both past and future projections (ranging from years 1600-2100) that can be used to model not only historical but also future changes in climate suitability (see Section 3.1 of the main text for an example of this).

ERA5 and ISIMIP use different units of precipitation (mm/s in ISIMIP and m/day in ERA5), we, thus, transformed both to a standard unit of mm/day. Then, since the daily precipitation is usually 0 in most locations on most days, and the timescale of change in precipitation is unlikely to affect species at a daily scale, we aggregated precipitation data at the monthly scale instead of the daily using the `monsum` function from cdo. We also aggregated this into a yearly precipitation value using the `yearsum` function from cdo.

To avoid the size differences between grid-cells of equal latitudinal extent, we reprojected the climate rasters of daily tasmax, daily tasmin, monthly precipitation, and yearly precipitation from each climate record into 24x24km grid-cells in EPSG:6933 (Brodzik et al., 2012) using bilinear interpolation (`remapbil` function from cdo). We chose a 24 km resolution to achieve a similar value to the original ERA5 resolution of roughly 31km (0.25°). The resolution of ISIMIP models differ depending on the specific model, but we selected a standard resolution of 24km for these as well to have synergy with ERA5. Despite potential confounding factors arising from topographic constraints and microclimatic buffering (Colwell, 2021), a 24kmx24km resolution can likely capture ecological patterns adequately (Hurlbert and Jetz, 2007; Trisos et al., 2020). Then, since the rasters are provided in temporal blocks representing a set of years, we merged each block's raster data (using `mergetime` from cdo) for the daily tasmax, daily tasmin, monthly precipitation, and yearly precipitation into one complete raster representing time-series for each variable over the whole historical period.

### S2 Local Climate Limits

From the climate rasters, we then computed the local climate limits as 44 metrics indicating the most acute temperature/precipitation conditions experienced in each of the 875,520 grid-cells over the relevant historical period. In particular, these local climate limits belong to three aspects- (a) maxima/minima in temperature and precipitation based on percentiles (26 local climate limits), (b) maximum annual variability of temperature and precipitation (4 local climate limits), and (c) extreme events in temperature and precipitation, quantified as their frequency, intensity, duration, and severity (14 local climate limits). In the below sections, we detail the computation of these metrics and their potential ecological relevance.

#### S2.1 Maxima/minima in temperature/precipitation

Maxima/minima limits in temperature and precipitation represent the most acute conditions experienced in a focal grid-cell over the historical period, and indicate the values that represent boundaries of expected climate conditions in that grid-cell. We computed these limits from climate records using different percentiles of 95, 99, 99.9 and 100 (and correspondingly 5, 1, 0.1 and 0 for minima) based on the common use of these percentile values in modeling extreme values. The reason we do this is to present a range of threshold conditions. This is to potentially deal with outliers being produced in the generation of these global climate records. This is in alignment with other studies of climate extremes (Murali et al., 2023). Another benefit is providing different cutoff values for the limits depending on the sensitivity of the metrics in assessing climate vulnerability for a focal species (for example, a more conservative prediction of future climate exposure may consider the 95th percentile being breached as an indication of stressful conditions in the grid-cell, rather than the 99th percentile). We computed these using `timselpctl` in cdo for each climate variable's raster in each model.

In addition to computing percentiles based on the whole time-series, we also computed a threshold based on seasonally varying climatology values. A climatology in this context refers to a curve indicating the value of a specified percentile of temperature values from all years in a time-window around a specified day. For example, one data point in a 95th percentile tasmax climatology would be the 95th percentile of the average tasmax values from a window of 5 days past and future (for a total of 10 days) around March 15 across each year in a time-series. We reported a 'seasonal maximum temperature' and 'seasonal minimum temperature' based on the highest/lowest value on the climatology curve, thus representing the most extreme temperature in the warmest/coldest period of the year. This tends to display more extreme values than the approach of computing the percentile over the whole time-series, since the whole time-series includes colder (/warmer) months in computing the percentile, which brings down (/up) the inferred limit.

We also highlight that we computed both the monthly and annual precipitation since these represent different properties - the monthly precipitation maximum represents shorter-term extremes, while annual precipitation reflects conditions including wet and dry seasons and therefore indicates more large-scale changes in the annual precipitation cycle. In addition, we computed the median precipitation in the wettest and driest month from each year using the `yearmin` and `yearmax` cdo functions and then taking the median. This provides further insight into local extremes, by comparing only between the maxima/minima of all years and not including the other seasons, thereby showcasing a greater range of conditions. We used the median to indicate what an expected amount of precipitation would be in the most extreme seasonal period.

#### S2.2 Annual variability

Climate variability is generally regarded as a major driver of ecological vulnerability and organismal limits (Janzen, 1967). Specifically, species which experience more variable environmental conditions historically are generally regarded as having lesser sensitivity to future extreme conditions (Sunday et al., 2011; Khaliq et al., 2014). Therefore, accounting for climatic variability in structuring climate limits enables us to model not only long-term changes in the average conditions and their limits, but also the likelihood of observing extreme values. This is especially pertinent given the contrasting trends of future change in climate variability around the planet (Bathiany et al., 2018).

We quantified annual variability in daily temperature and monthly precipitation conditions in a grid-cell as the coefficient of variation and the standard deviation of conditions over the year. For temperature, since we use both the daily maximum and minimum temperature, we computed the variability of the average daily temperature by taking the average value of the tasmax and tasmin for each day. By returning the value of the coefficient of variation and standard deviation for each year for each grid-cell, we computed the local climate limit for the grid-cell as the 99th percentile of the variability values from each year (with a similar motive as using the percentile, to remove any potential outlier values produced by the model). This was done using the `yearmean` and `yearstd1` functions from cdo.

#### S2.3 Extreme Events

Extreme events represent sustained periods of extreme supra-threshold conditions in temperature and precipitation, in the form of heatwaves, cold-spells, and droughts.

We defined two types of extreme temperature events, the computation of which is summarized in panels (a) and (b) of Figure S1. We termed these as 'heatwaves' and 'maximum heatwaves' (correspondingly, 'cold-spells' and 'minimum cold-spells'). For modeling both types of events, we used the approach of constructing a climatology as a curve representing the value of a given percentile of temperature values from all years in a 10-day time-window around a specified day (represented by the dotted line in Figure S1 (a) and (b)). In this framework, the first type of events, heatwaves (and cold-spells) are defined as periods when the temperature exceeds the value of the climatological curve for five consecutive days or more (Perkins and Alexander, 2013; Hobday et al., 2016) (as in Figure S1 (a)). This stipulates heatwaves (or cold-spells) as periods when conditions are warmer (or colder) than the conditions typically experienced in this time of year for a sustained duration. Specifically, for heatwaves (and cold-spells), we fixed the percentile of the climatology as the 99th percentile (or 1st percentile), and computed periods where the tasmax (or tasmin) is above (or below) the climatology. We then computed the frequency of heatwave and cold-spell events over the historical period as the average number of events per year.

Heatwaves and cold-spells are understood to have drastic effects on ecosystems by affecting life history, interactions, behavior, and phenology (Regan and Sheldon, 2023; Martínez-De León and Thakur, 2024). For example, a heatwave in the winter may significantly affect ecological processes by causing early snowmelt. However, these events outside the warmest period may not lead to significant physiological stress outside the thermal limits of an organism exposed to significantly higher summer temperatures (Broadbent et al., 2024). Therefore, we defined the second type of extreme temperature events, 'maximum heatwaves' (or 'minimum cold-spells') as periods which may indicate significant thermal stress for species by occurring in the warmest (or coldest) period of the year (as in Figure S1 (b)). To model maximum heatwaves (and minimum cold-spells), we computed the maximum value of the annual climatology (similar to the computation of seasonal maximum temperature described in Section S2.1), and used this as a threshold temperature for the organism. In this framework, a maximum heatwave (or minimum cold-spell) is a period of five days or more where the temperature conditions exceed the global maximum (or minimum) of the climatology. This indicates conditions more extreme than the maximum value across all time-windows, and may have severe effects on an organism, including necrotic tissue damage, dehydration, and starvation (Ma et al., 2015; Till et al., 2019; Román-Palacios and Wiens, 2020). By construction, maximum heatwave (or minimum cold-spell) events are typically biased towards the warmest summer months (or coldest winter months). We fixed the 99th/1st percentiles for the climatology to define the threshold and measure these events.

For maximum heatwaves and minimum cold-spells, we computed the frequency as the average number of events per year, but additionally, for each grid-cell, we computed the maximum duration as the longest event recorded in the grid-cell (in terms of number of days), the maximum intensity of all events observed across the grid-cell (measured for each event as the mean difference between the observed temperature and the threshold defined by the maximum/minimum of the climatology during the event), and the maximum severity across all events in the grid-cell (defined for each event as the duration multiplied by the intensity). The severity is also included on account of the negative correlation between duration and intensity as a consequence of their definitions. We did not compute these properties for the heatwaves/cold-spells (the first type of extreme temperature events) since these represent events that could happen at any time of year, and thus their effects may be less direct on organisms.

Climatologies were constructed using the `ts2clm` function from the 'heatwaveR' package (version 0.5.4) in R (Schlegel and Smit, 2018). Heatwaves and cold-spells were computed from these climatologies using the `detect_event` function from 'heatwaveR' - maximum heatwaves were detected by extracting the threshold from the climatology and using a custom function to detect periods of exceedance.

For extreme events in precipitation, we computed droughts as periods of reduced precipitation. Though our framework may also be feasible for computing floods as periods of elevated precipitation, we did not do this since the spatial scale of a flooding event is typically smaller than the 24km x 24km resolution we use (de Moel et al., 2015; Savage et al., 2016).

For detecting drought, we computed the SPEI (Standardized Precipitation Evapotranspiration Index) (Vicente-Serrano et al., 2010). The SPEI is a standard approach to determine the onset, duration, and magnitude of drought events. It is computed as the inverse of a cumulative distribution function for a value of given rainfall for a given period of a year. This distribution function in turn is computed based on standardizing the water deficit for each period, defined as the difference between the observed precipitation and the potential evapotranspiration (PET). We measured the PET for each period using the Hargreaves method (Hargreaves and Samani, 1985), which uses the temperature, observed rainfall, and latitude of a grid-cell to compute the expected evapotranspiration for that grid-cell. We computed the SPEI at a timescale of 3 months - this was selected based on its relevance to ecological timescales, as opposed to longer-term patterns of hydrological or vegetative changes. (Crausbay et al., 2017). With this, we defined a drought event according to standard criteria (Medri et al., 2020), which stipulate a drought beginning when the SPEI is less than -1 for two consecutive periods, and ends when the SPEI becomes positive, indicating that the period of decreased rainfall has ended (see Figure S1 (c)). This records a drought event at a monthly scale.

We calculated the SPEI values using the `spei` and `hargreaves` functions from the R package 'SPEI' (version 1.8.1) (Beguería and Vicente-Serrano, 2023). For each grid-cell, we measured the frequency of drought events as the average number of drought events per year. In addition, we also measured the maximum duration as the longest drought event (in months) in the grid-cell, the maximum intensity of all drought events in the grid-cell (quantifying intensity as the mean SPEI during the event), and maximum severity of all drought events in the grid-cell (quantifying severity as the duration of the event multiplied by its intensity). For some grid-cells with strong dry seasons, the SPEI can occasionally produce values of -Inf due to failure to fit values to the distribution successfully - we did not compute drought events in these grid-cells and excluded them from the analysis. As a consequence of this, for 121 species whose ranges were totally comprised of such grid-cells, it was not possible to compute species climate limits - for these species, we do not compute the species climate limits of drought duration, intensity, and severity, and report these as NA in the final output.

### S3 Species Climate Limits

We estimated species climate limits from local climate limits at the grid-cell level based on the annotated species' geographic range. This was done for each local climate limit by identifying the grid-cells where the species was found, extracting the value of the local climate limit in each of the grid cells, and taking the maximum or minimum over all local climate limit values (computed as per Section S2). This produced a single realized species-level limit from each local climate limit, producing a final set of 44 species climate limits for each species (listed out in Table 1 in main text). For example, the species climate limit $T_{max99}$ is the maximum value over all grid-cells in the species' range of the local climate limit of 99th percentile of daily maximum temperature; correspondingly, the species climate limit $T_{min1}$ is the minimum value of the local climate limit of the 1st percentile of daily minimum temperature over the species' geographic range. For extreme events, we took the species' maximum duration, intensity, and severity similarly by comparing events across all grid-cells constituting the species' range.

#### S3.1 Correlations between species climate limits

Many of the computed species climate limits are variations on a single threshold, computed at different limiting percentiles- for example, we compute the maximum temperature and maximum monthly rainfall at the 99th, 99.9th, and 100th percentiles. This is done intentionally - we provide thresholds depending on

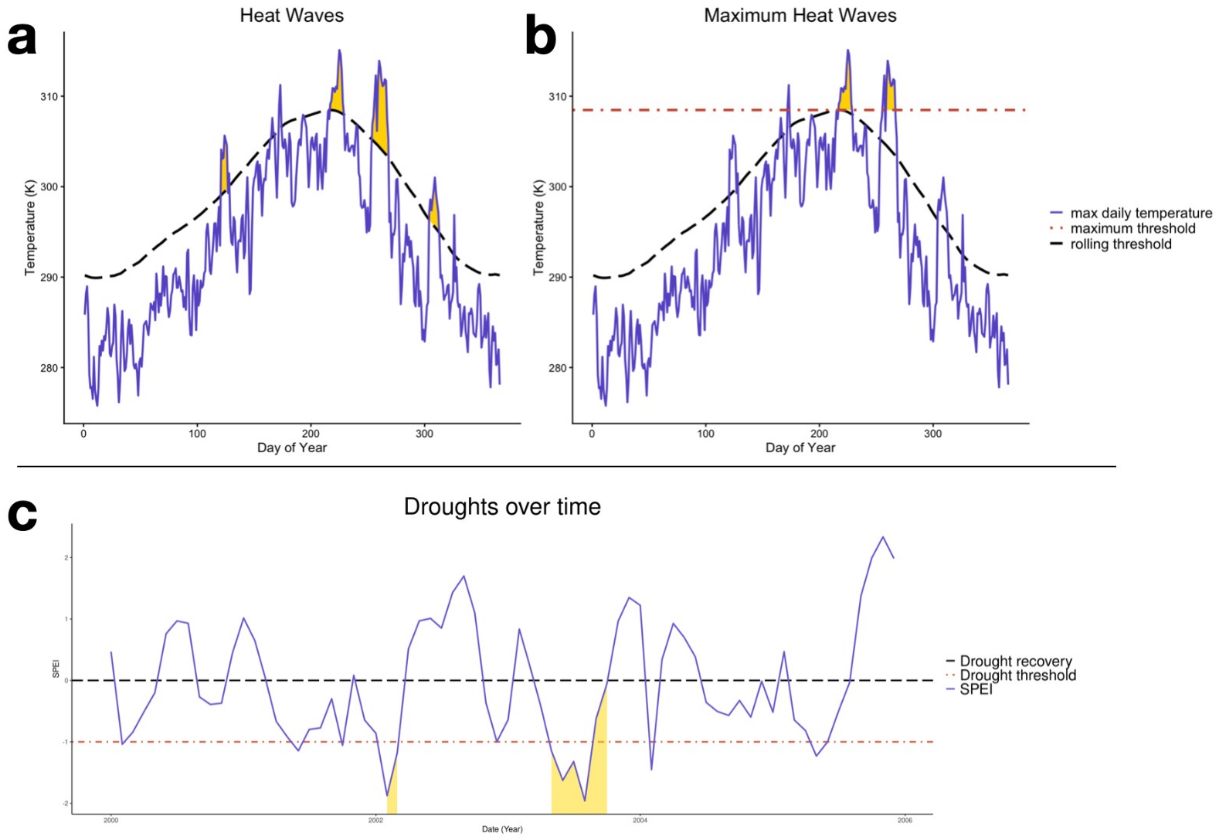


Figure S1: **Computation of extreme events in temperature and rainfall.** (a) This panel depicts the computation of heatwaves based on the annual climatology. The climatology, constructed as a percentile of values of temperature around a time-window for each day of the year, is depicted as a dotted line. The sample time-series is depicted in blue, with the heatwave events as sustained periods of exceedance of the climatological limit highlighted in yellow. (b) This panel depicts the computation of maximum heatwaves using the climatological approach. The same time-series of climate from (a) is shown in blue, with maximum heatwaves above the threshold line (in red) highlighted in yellow. Note that the heatwaves shown in panel (a) have different occurrence and properties than the maximum heatwaves. (c) This panel shows a sample time-series of SPEI, and the computation of drought from this. A drought begins when the SPEI is less than -1 for two consecutive months, and ends when the SPEI value becomes positive.

how conservatively the user plans to assess the climate limits of a focal species. Many of the variables are also inherently correlated (for example, the duration and intensity of maximum heatwaves are negatively correlated, since the probability to sustain an intensely high temperature for a long period is lower). Figures S2, S3, S4, S5, S6, and S7 show the correlation between all 44 climate limits across all 54,255 species in the ClimLimits database, for each of the climate records. We note that patterns of correlation are largely consistent between different climate records. When modeling niche dimensions for a focal species, we recommend considering these correlations to select an optimal set of variables.

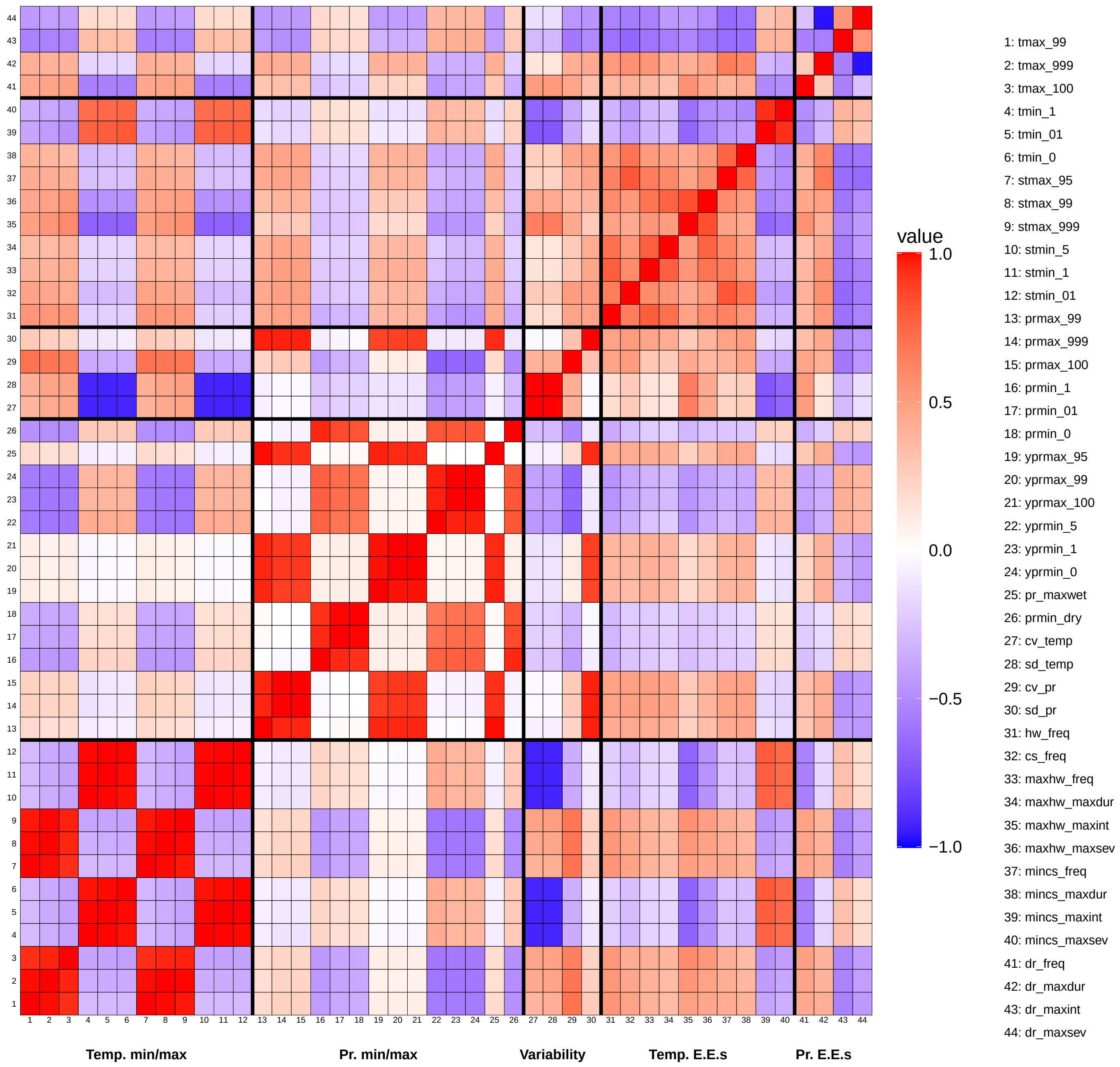


Figure S2: **Correlogram for ERA5**. This indicates strong patterns of correlation within aspects, as well as some correlations between variables in different aspects.

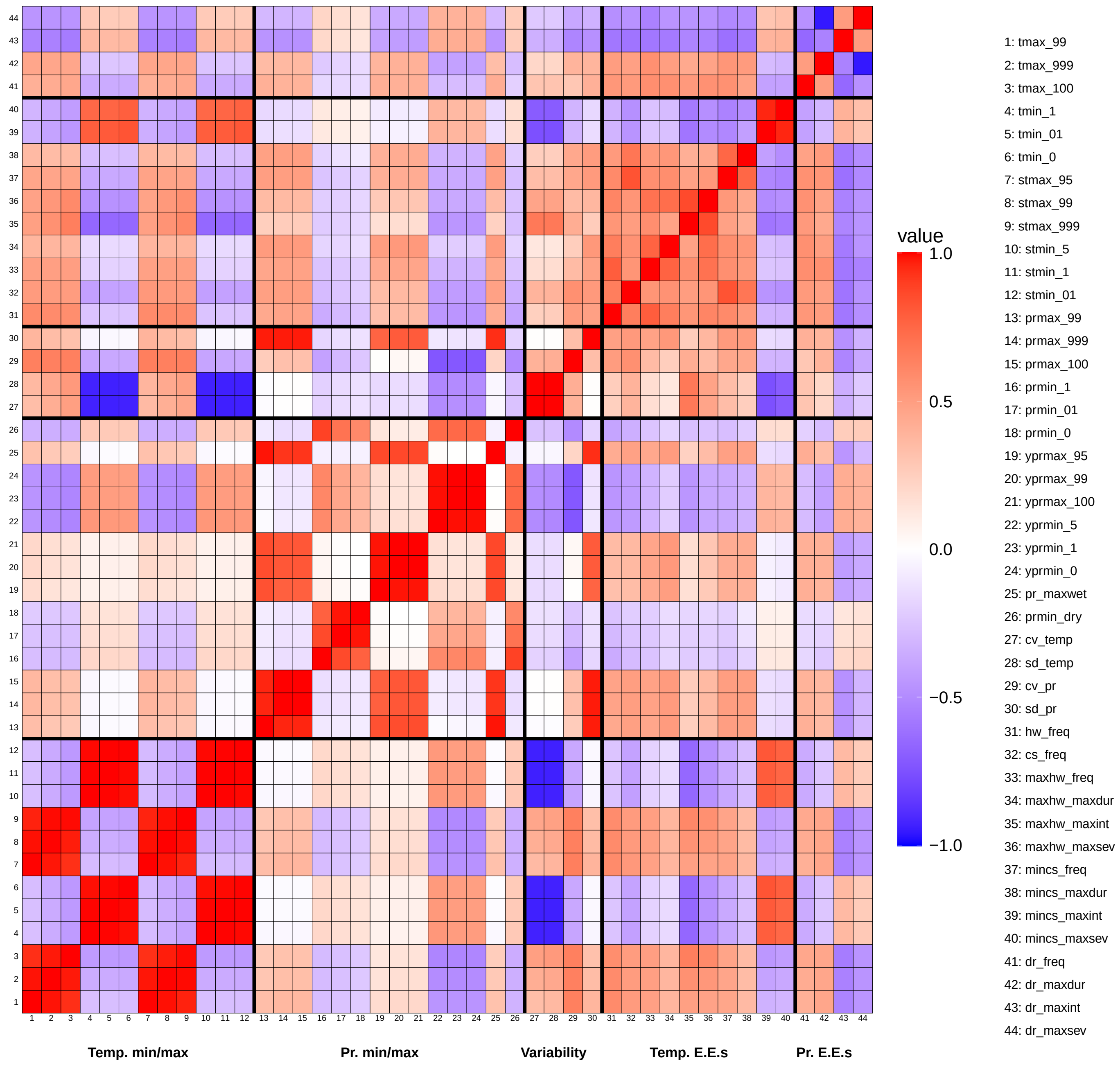


Figure S3: **Correlogram for EC-EARTH3**. This indicates strong patterns of correlation within aspects, as well as some correlations between variables in different aspects.

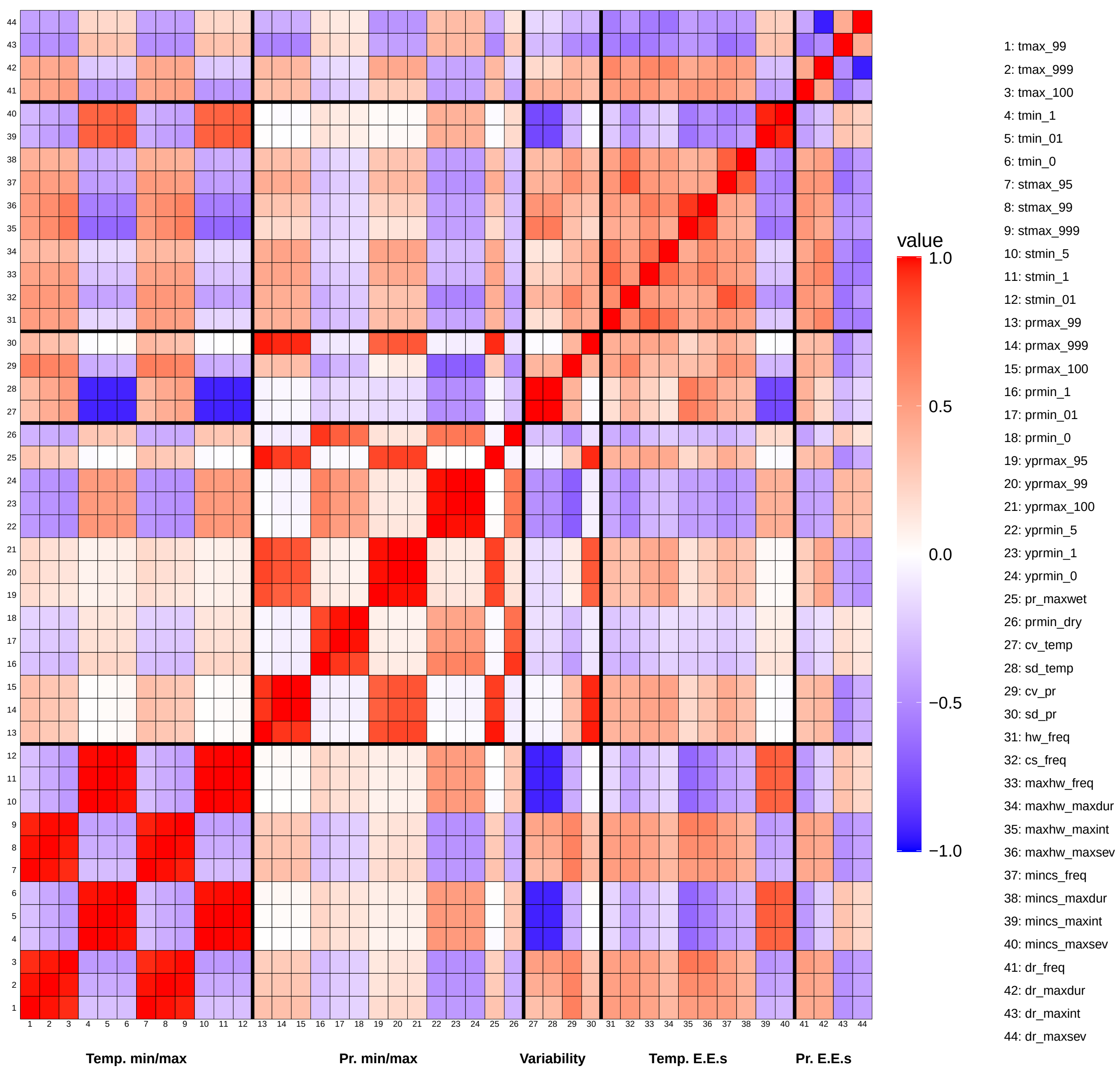


Figure S4: **Correlogram for IPSL-CM6A-LR**. This indicates strong patterns of correlation within aspects, as well as some correlations between variables in different aspects.

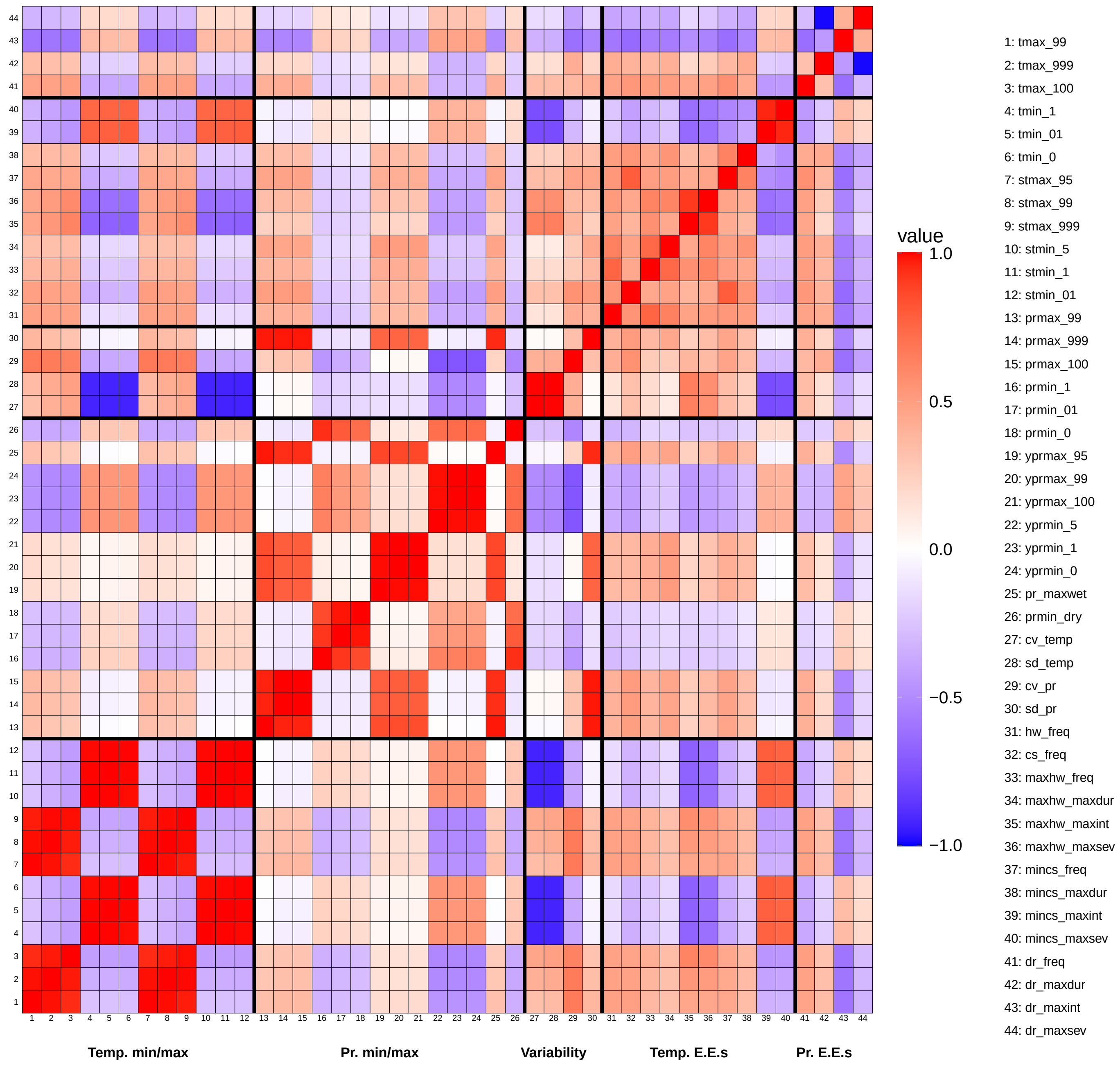


Figure S5: **Correlogram for GFDL-ESM4**. This indicates strong patterns of correlation within aspects, as well as some correlations between variables in different aspects.

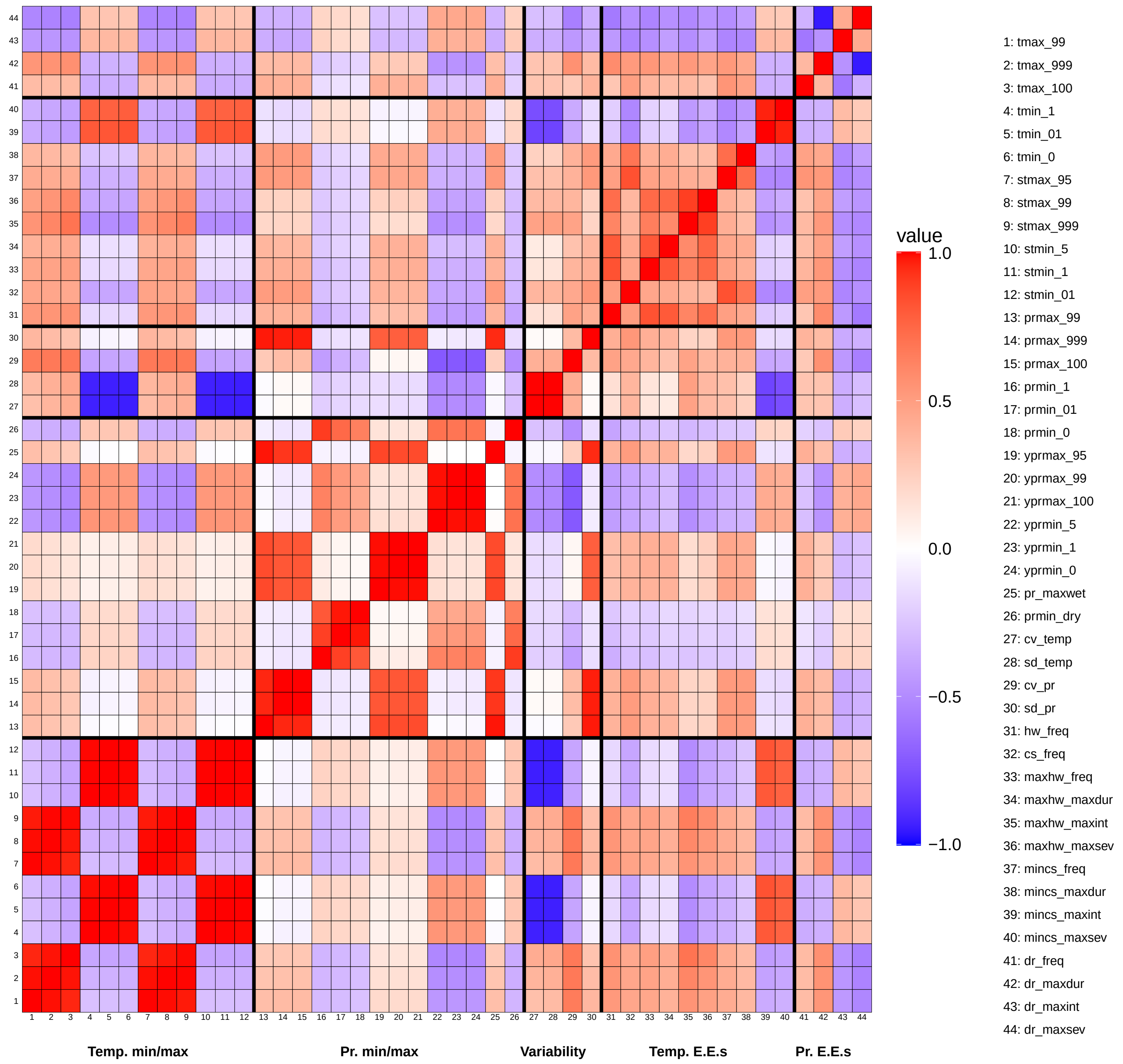


Figure S6: **Correlogram for MIROC6**. This indicates strong patterns of correlation within aspects, as well as some correlations between variables in different aspects.

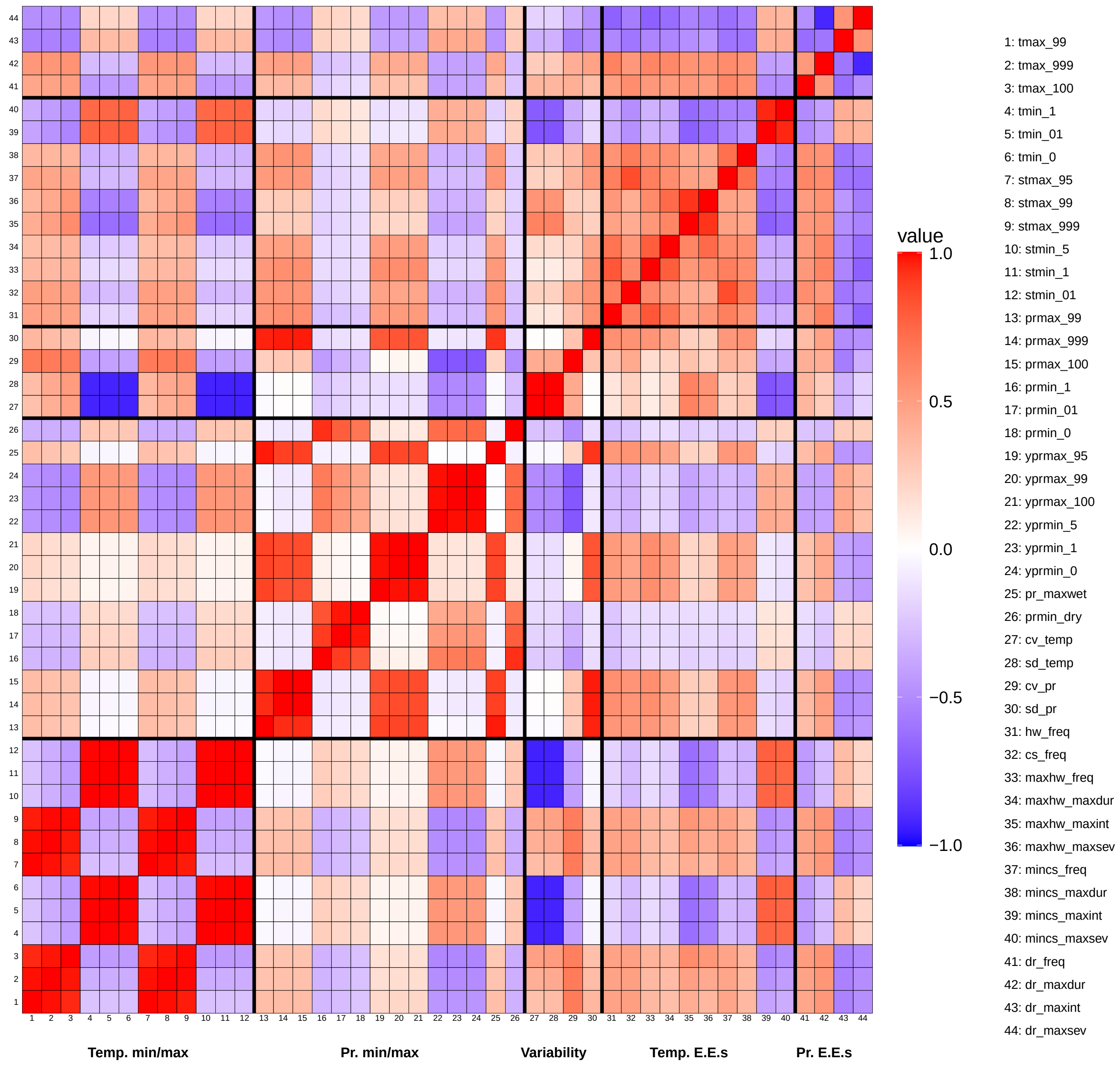


Figure S7: **Correlogram for MPI-ESM1-2-HR**. This indicates strong patterns of correlation within aspects, as well as some correlations between variables in different aspects.

#### S3.2 Species climate limits on taxonomic and geographical subsets

We recognize that the spatial scale of our database is extensive, and requires processing a tremendous amount of species range and climate record data. Indeed, this is one of the major motivations for producing the ClimLimits database in this format, to provide an easy-to-access pre-computed global record of realized species climate limits for climate change vulnerability assessments. However, for reproducing the database on smaller geographic scales for regional-scale assessments, we also provide a "subset" functionality with our script. This considers only a smaller pool of the total species, as well as a smaller geographic area, making local-scale estimates of species climate limits that also require less computing. This facilitates the production of regional-scale assessments of species climate limits focusing only on a focal area or group of taxa. Regional-scale species climate limits may be different from global-scale climate limits, revealing local patterns of sensitivity.

As a sample, along with the code to produce the database, we provide pre-processed climate data from the ERA5 model on daily maximum/minimum temperature and monthly/annual precipitation for a bounding box from 41N to 51N in latitude, and -6E to 10E in longitude (covering the region of Metropolitan France and some other parts of the European continent around it) along with the database as an additional folder. The script to generate species climate limits can be run on a specified subset by providing a run type ('full' or 'subset') and a subset code (the one we have provided is called 'metrofrance') in the script. The species groups for which species climate limits will be generated can also be specified in the script - for the provided sample, we annotate species climate limits for all mammals in this bounding box (representing 113 species/177 populations). We also provide a script to generate a regional-level subset from global climate records (`climlimits_08_subset_generator.R`) along with all required peripheral files - by selecting latitude/longitude bounds and specifying a subset name, the user may then generate species climate limits for this region for a specified group of taxa.

### S4 Species Range Visualization

Another functionality we provide in the ClimLimits database is the option to visualize the ranges of species and their constituent populations on a map, using the provided additional data of the annotated species' range, and the boundaries of each grid-cell. This tool is provided in the script `climlimits_05_case_study.R`, which defines a function `visualize_range` to use the range data and grid-cell information to produce a map of the species range. This function provides several options for producing the map, specifically- (a) pop_struct depicts each population of the species separately if it is ON, and groups them into one common range if it is OFF; (b) cells as an argument allows one to visualize the grid-cells and the shapefile constituting the range if it is ON, and only the original shapefile if it is OFF; (c) the world_map argument plots the range on the background of a world map if it is ON, and plots only the range if OFF; and (d) the pop_number argument plots all populations if it is specified as NA (default), and a specific population number if provided with one. We note that the population numbers are assigned arbitrarily - thus, if interested in the range of a single focal population, we recommend running the function with pop_struct set to ON and pop_number set to NA, identifying the population of interest on the map, then rerunning the function with only that number specified as argument of pop_number. Figure S8 depicts a sample output from this function from a species with 3 populations, the Tantalus Monkey (*Chlorocebus tantalus*). This was done with the options of pop_struct ON, cells OFF, world_map ON, and pop_number NA, to visualize all populations' original shapefiles- specifically, the function call done is `visualize_range(sp_name="Chlorocebus tantalus", pop_struct="ON", cells="OFF", world_map="ON", pop_number=NA)`.

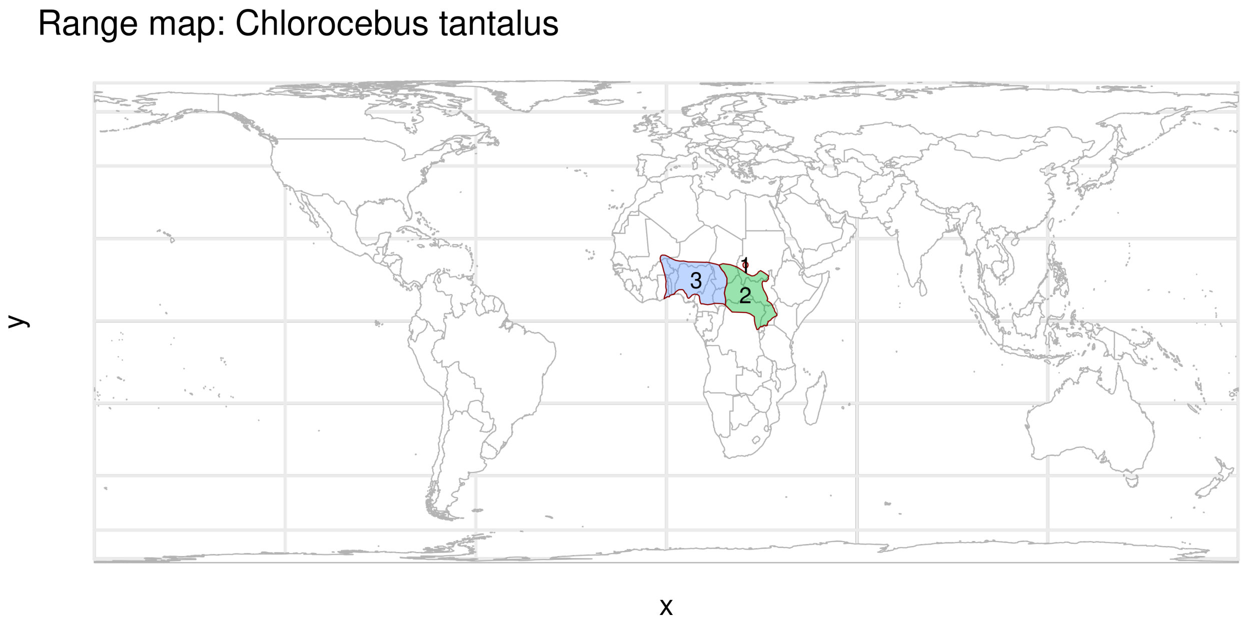


Figure S8: **Sample range map output from the `visualize_range` function**, for the given species Tantalus Monkey (*Chlorocebus tantalus*). The species has 3 distinct populations annotated by the IUCN - this map was produced by provoding the arguments pop_struct ON, cells OFF, world_map ON, and pop_number NA, thereby producing a visualization of all shapefiles for the species' populations on the world map.

#### S5 Species Climate Limits and Range Shifts

One bias that could impact our estimates of realized species climate limits over their range is that in our analysis, we use a single static snapshot of the species' range annotated by the IUCN, compared against dynamic records of climate conditions experienced over the last 80 years in their current range. However, a wealth of evidence has shown that species are currently shifting their geographic ranges in response to climate change (Parmesan and Yohe, 2003; Lenoir and Svenning, 2014; Rubenstein et al., 2023); indeed, these ranges could be moving over the reference period for climates, meaning that a species may have been found in a particular cell in the past but not the present (or vice versa), affecting the estimates of species climate limits that we make. To quantify this the scale of this effect, we used the BioShifts database (Lenoir et al., 2020), which provides a synthesis of estimated annual shift rates for many species across both latitude and elevation. These estimated shift rates, based often on fairly recent sampling efforts (compared to the historical span of our database), may be slightly higher than the past on account of the acceleration of climate warming in recent decades and the ensuing need for species to track suitable climate conditions in space. Therefore, we can consider this as an upper bound on how much our estimated ranges could shift in an 80-year historical window (assuming there is no changes in landscape and habitat connectivity level constraints throughout this range shifting period).

We select all records from the BioShifts database of animals undergoing latitudinal range shifts in terrestrial ecosystems, since errors in our estimation of species climate limits will largely emerge from latitudinal range shifts. We then subset to only taxa in the ClimLimits database (freshwater fish, amphibians, birds, crabs, mammals, and reptiles). Though these rates have varying qualities, with some being modeled as opposed to being resurveyed empirically, or may quantify shifts in the centroid or either of the range edges which could have different rates, we retain all of these to have as much species coverage as possible. Of 1,354 species in the BioShifts database with terrestrial latitudinal shifts, we subset to only those present in the ClimLimits database. This provides us with 4,484 distinct values of shift rate for 1,111 species from 51 studies. For species with multiple estimates of range shift rate, we then average them into a single rate per species (in km/year). Then, we can transform this into a number of cells the species is likely to have moved in 80 years by multiplying the rate by 80 (the number of years in the ERA5 dataset) and then dividing by 24 (the length of a single grid-cell).

Figure S9a depicts a violin plot of the number of cells species are likely to move in this period assuming they undergo current shift rates throughout the historical period. We see that while there are some outlier species undergoing high amounts of shift upto 112 cells, the median is 3.66 cells, with 85.5% of species (951 out of 1,111) experiencing shifts of less than 10 cells. Note that a large majority of these species analyzed here are birds (roughly 84% of the 1,111 species), which are likely more motile than many other species (for example, amphibians).

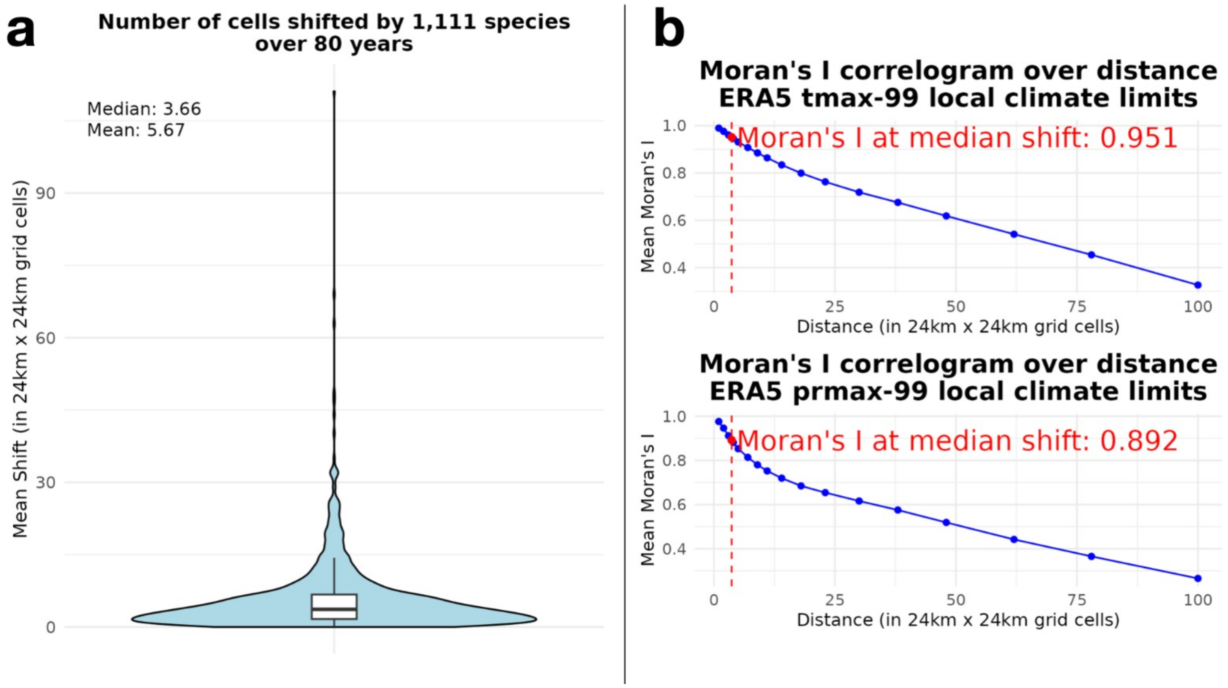


Figure S9: **Quantifying the effect of species range shifts on species climate limits estimates**. Panel (a) shows, given the rates of shift estimated per year from the BioShifts database for 1,111 species, the number of 24km x 24km grid-cells species are expected to shift latitudinally during the 80-year historical baseline period. We see a median shift of 3.66 cells. Panel (b) quantifies how much average correlation there is between local climate limit values over space by quantifying average Moran's I coefficients as a function of distance around each grid-cell. This is shown for the local climate limits of tmax-99 (99th percentile of maximum daily temperature) and prmax-99 (99th percentile of maximum monthly precipitation), and shows a high degree of correlation for most smaller values of shift - notably, for the median shift amount, we get coefficients of 0.95 and 0.89 for tmax-99 and prmax-99 respectively. This shows that correlations remain high, indicating that range shifts may impact estimates, but the scale of change is limited.

To quantify how much shift this actually means in a climatic sense, we find the spatial autocorrelation of different local climate limits. This measures how similar local climate limits are present in windows of different cell-sizes around a focal cell. By picking cells a specified distance away from each grid-cell and computing their Moran's I-coefficients, then averaging these values across all grid-cells, we can find how much spatial autocorrelation is expected for a climate limit given a distance. This quantifies how similar these limiting values are across grid-cells. Figure S9b depicts these plots for two sample local climate limits - 99th percentile of temperature (tmax-99) and 99th percentile of monthly precipitation (prmax-99). This indicates that at small numbers of grid-cells, the spatial autocorrelation is quite high indicating a small degree of change - indeed, at the median value of shift in the dataset (3.66 cells), we see the Moran's I coefficient is 0.95 for temperature and 0.89 for precipitation, indicating that these values are very well-correlated to the estimated limits. For the species climate limits, since these are computed as the global maximum/minimum over local climate limits stretching over the species' range, these are likely to indicate even higher correlation, since the distribution changes may not affect the extreme values over all the grid-cells in the species' range. Thus, while range shifts can indeed impact species distributions over the study period, they are unlikely to modify the species' climate limits we have estimated to an appreciable extent.

### References

Alexander, J. M., Chalmandrier, L., Lenoir, J., Burgess, T. I., Essl, F., Haider, S., Kueffer, C., McDougall, K., Milbau, A., Nuñez, M. A., Pauchard, A., Rabitsch, W., Rew, L. J., Sanders, N. J., and Pellissier, L. (2018). Lags in the response of mountain plant communities to climate change. *Glob. Chang. Biol.*, 24(2):563–579.

Bathiany, S., Dakos, V., Scheffer, M., and Lenton, T. M. (2018). Climate models predict increasing temperature variability in poor countries. *Sci. Adv.*, 4(5):eaar5809.

Beguería, S. and Vicente-Serrano, S. M. (2023). *SPEI: Calculation of the Standardized Precipitation-Evapotranspiration Index*. https://spei.csic.es, https://github.com/sbegueria/SPEI.

Birdlife, I. (2024). Birdlife — partnership for nature and people.

Boonman, C. C. F., Hoeks, S., Serra-Diaz, J. M., Guo, W.-Y., Enquist, B. J., Maitner, B., Merow, C., and Svenning, J.-C. (2025). High tree diversity exposed to unprecedented macroclimatic conditions even under minimal anthropogenic climate change. *Proceedings of the National Academy of Sciences*, 122(26).

Broadbent, A. A., Newbold, L. K., Pritchard, W. J., Michas, A., Goodall, T., Cordero, I., Giunta, A., Snell, H. S., Pepper, V. V., Grant, H. K., et al. (2024). Climate change disrupts the seasonal coupling of plant and soil microbial nutrient cycling in an alpine ecosystem. *Global Change Biology*, 30(3):e17245.

Brodzik, M. J., Billingsley, B., Haran, T., Raup, B., and Savoie, M. H. (2012). EASE-Grid 2.0: Incremental but Significant Improvements for Earth-Gridded Data Sets. *ISPRS International Journal of Geo-Information*, 1(1):32–45.

Chamberlain, S., Barve, V., Mcglinn, D., Oldoni, D., Desmet, P., Geffert, L., and Ram, K. (2026). *rgbif: Interface to the Global Biodiversity Information Facility API*. R package version 3.8.5.

Chuine, I. (2010). Why does phenology drive species distribution? *Philosophical Transactions of the Royal Society B: Biological Sciences*, 365(1555):3149–3160.

Colwell, R. K. (2021). Spatial scale and the synchrony of ecological disruption. *Nature*, 599(7886):E8–E10.

Crausbay, S. D., Ramirez, A. R., Carter, S. L., Cross, M. S., Hall, K. R., Bathke, D. J., Betancourt, J. L., Colt, S., Cravens, A. E., Dalton, M. S., Dunham, J. B., Hay, L. E., Hayes, M. J., McEvoy, J., McNutt, C. A., Moritz, M. A., Nislow, K. H., Raheem, N., and Sanford, T. (2017). Defining ecological drought for the twenty-first century. *Bull. Am. Meteorol. Soc.*, 98(12):2543–2550.

de Moel, H., Jongman, B., Kreibich, H., Merz, B., Penning-Rowsell, E., and Ward, P. J. (2015). Flood risk assessments at different spatial scales. *Mitig. Adapt. Strateg. Glob. Chang.*, 20(6):865–890.

Eyring, V., Cox, P. M., Flato, G. M., Gleckler, P. J., Abramowitz, G., Caldwell, P., Collins, W. D., Gier, B. K., Hall, A. D., Hoffman, F. M., Hurtt, G. C., Jahn, A., Jones, C. D., Klein, S. A., Krasting, J. P., Kwiatkowski, L., Lorenz, R., Maloney, E., Meehl, G. A., Pendergrass, A. G., Pincus, R., Ruane, A. C., Russell, J. L., Sanderson, B. M., Santer, B. D., Sherwood, S. C., Simpson, I. R., Stouffer, R. J., and Williamson, M. S. (2019). Taking climate model evaluation to the next level. *Nature Climate Change*, 9(2):102–110.

Flato, G., Marotzke, J., Abiodun, B., Braconnot, P., Chou, S. C., Collins, W., Cox, P., Driouech, F., Emori, S., Eyring, V., et al. (2014). Evaluation of climate models. In *Climate change 2013: the physical science basis. Contribution of Working Group I to the Fifth Assessment Report of the Intergovernmental Panel on Climate Change*, pages 741–866. Cambridge University Press.

Fu, Z., Ciais, P., Feldman, A. F., Gentine, P., Makowski, D., Prentice, I. C., Stoy, P. C., Bastos, A., and Wigneron, J.-P. (2022). Critical soil moisture thresholds of plant water stress in terrestrial ecosystems. *Science Advances*, 8(44).

Garcia, R. A., Burgess, N. D., Cabeza, M., Rahbek, C., and Araújo, M. B. (2012). Exploring consensus in 21st century projections of climatically suitable areas for african vertebrates. *Global Change Biology*, 18(4):1253–1269.

GBIF (2025). GBIF — gbif.org. `https://www.gbif.org`. [Accessed 11-12-2025].

Grossiord, C., Buckley, T. N., Cernusak, L. A., Novick, K. A., Poulter, B., Siegwolf, R. T. W., Sperry, J. S., and McDowell, N. G. (2020). Plant responses to rising vapor pressure deficit. *New Phytologist*, 226(6):1550–1566.

Hargreaves, G. H. and Samani, Z. A. (1985). Reference crop evapotranspiration from temperature. *Applied Engineering in Agriculture*, 1(2):96–99.

Hawkins, E. and Sutton, R. (2011). The potential to narrow uncertainty in projections of regional precipitation change. *Climate Dynamics*, 37(1-2):407–418.

Hersbach, H., Bell, B., Berrisford, P., Biavati, G., Horányi, A., Muñoz Sabater, J., Nicolas, J., Peubey, C., Radu, R., Rozum, I., Schepers, D., Simmons, A., Soci, C., Dee, D., and Thépaut, J.-N. (2023). Era5 hourly data on single levels from 1940 to present.

Hobday, A. J., Alexander, L. V., Perkins, S. E., Smale, D. A., Straub, S. C., Oliver, E. C., Benthuysen, J. A., Burrows, M. T., Donat, M. G., Feng, M., Holbrook, N. J., Moore, P. J., Scannell, H. A., Sen Gupta, A., and Wernberg, T. (2016). A hierarchical approach to defining marine heatwaves. *Progress in Oceanography*, 141:227–238.

Hurlbert, A. H. and Jetz, W. (2007). Species richness, hotspots, and the scale dependence of range maps in ecology and conservation. *Proceedings of the National Academy of Sciences*, 104(33):13384–13389.

IUCN (2025). The iucn red list of threatened species.

Janzen, D. H. (1967). Why Mountain Passes are Higher in the Tropics. *The American Naturalist*, 101(919):233–249.

Karl, T. R., Nicholls, N., and Ghazi, A. (1999). Clivar/gcos/wmo workshop on indices and indicators for climate extremes workshop summary. *Climatic Change*, 42(1):3–7.

Khaliq, I., Hof, C., Prinzinger, R., Böhning-Gaese, K., and Pfenninger, M. (2014). Global variation in thermal tolerances and vulnerability of endotherms to climate change. *Proc. Biol. Sci.*, 281(1789):20141097.

Lange, S. (2019). Trend-preserving bias adjustment and statistical downscaling with ISIMIP3BASD (v1.0). *Geoscientific Model Development*, 12(7):3055–3070.

Lenoir, J., Bertrand, R., Comte, L., Bourgeaud, L., Hattab, T., Murienne, J., and Grenouillet, G. (2020). Species better track climate warming in the oceans than on land. *Nature ecology & evolution*, 4(8):1044–1059.

Lenoir, J. and Svenning, J. (2014). Climate-related range shifts – a global multidimensional synthesis and new research directions. *Ecography*, 38(1):15–28.

Ma, G., Hoffmann, A. A., and Ma, C.-S. (2015). Daily temperature extremes play an important role in predicting thermal effects. *Journal of Experimental Biology*, 218(14):2289–2296.

Martínez-De León, G. and Thakur, M. P. (2024). Ecological debts induced by heat extremes. *Trends in Ecology & Evolution*, 39(11):1024–1034.

Medri, S., Crespi, A., Terzi, S., Cocuccioni, S., Zebisch, M., Berckmans, J., and Füssel, H.-M. (2020). Climate-related hazard indices for europe. Technical Report Technical Paper 1/2020, European Topic Centre on Climate Change Adaptation (ETC/CCA), Bologna, Italy. Prepared by EURAC Research, VITO, and the European Environment Agency (EEA).

Murali, G., Iwamura, T., Meiri, S., and Roll, U. (2023). Future temperature extremes threaten land vertebrates. *Nature*, 615(7952):461–467.

Parmesan, C. and Yohe, G. (2003). A globally coherent fingerprint of climate change impacts across natural systems. *Nature*, 421(6918):37–42.

Pebesma, E. (2018). Simple Features for R: Standardized Support for Spatial Vector Data. *The R Journal*, 10(1):439.

Pebesma, E. and Bivand, R. (2023). *Spatial Data Science: With Applications in R*. Chapman and Hall/CRC, New York, 1 edition.

Perkins, S. E. and Alexander, L. V. (2013). On the Measurement of Heat Waves. *Journal of Climate*, 26(13):4500–4517.

Peterson, T. C. and Coauthors (2001). Report on the activities of the working group on climate change detection and related rapporteurs 1998-2001. wmo. *Rep.*, WCDMP-47, WMO-TD 1071, Geneve, Switzerland. 143pp.

Regan, C. E. and Sheldon, B. C. (2023). Phenotypic plasticity increases exposure to extreme climatic events that reduce individual fitness. *Global Change Biology*, 29(11):2968–2980.

Rheindt, F. E., Donald, P. F., Donsker, D. B., Gerbracht, J. A., Iliff, M. J., Lepage, D., Norman, J. A., Rasmussen, P. C., Schodde, R., Schulenberg, T. S., Areta, J. I., Brammer, F. P., Chesser, R. T., Dowsett, R. J., Peterson, A., Alström, P., Stervander, M., Remsen, J. V., Garnett, S. T., Homberger, D. G., Lei, F., and Christidis, L. (2025). Avilist: a unified global bird checklist. *Biodiversity and Conservation*, 34(10):3359–3376.

Román-Palacios, C. and Wiens, J. J. (2020). Recent responses to climate change reveal the drivers of species extinction and survival. *Proceedings of the National Academy of Sciences*, 117(8):4211–4217.

Roy, K., Jablonski, D., Valentine, J. W., and Rosenberg, G. (1998). Marine latitudinal diversity gradients: Tests of causal hypotheses. *Proceedings of the National Academy of Sciences*, 95(7):3699–3702.

Rubenstein, M. A., Weiskopf, S. R., Bertrand, R., Carter, S. L., Comte, L., Eaton, M. J., Johnson, C. G., Lenoir, J., Lynch, A. J., Miller, B. W., Morelli, T. L., Rodriguez, M. A., Terando, A., and Thompson, L. M. (2023). Climate change and the global redistribution of biodiversity: substantial variation in empirical support for expected range shifts. *Environmental Evidence*, 12(1).

Sandel, B., Merow, C., Serra-Diaz, J. M., and Svenning, J. (2025). Disequilibrium in plant distributions: Challenges and approaches for species distribution models. *Journal of Ecology*, 113(4):782–794.

Savage, J. T. S., Bates, P., Freer, J., Neal, J., and Aronica, G. (2016). When does spatial resolution become spurious in probabilistic flood inundation predictions? *Hydrol. Process.*, 30(13):2014–2032.

Schlegel, R. W. and Smit, A. J. (2018). heatwaveR: A central algorithm for the detection of heatwaves and cold-spells. *Journal of Open Source Software*, 3(27):821.

Schulzweida, U. (2023). Cdo user guide.

Sunday, J. M., Bates, A. E., and Dulvy, N. K. (2011). Global analysis of thermal tolerance and latitude in ectotherms. *Proc. Biol. Sci.*, 278(1713):1823–1830.

Svenning, J.-C. and Sandel, B. (2013). Disequilibrium vegetation dynamics under future climate change. *Am. J. Bot.*, 100(7):1266–1286.

Tebaldi, C. and Knutti, R. (2007). The use of the multi-model ensemble in probabilistic climate projections. *Philosophical Transactions of the Royal Society A: Mathematical, Physical and Engineering Sciences*, 365(1857):2053–2075.

Till, A., Rypel, A. L., Bray, A., and Fey, S. B. (2019). Fish die-offs are concurrent with thermal extremes in north temperate lakes. *Nature Climate Change*, 9(8):637–641.

Trisos, C. H., Merow, C., and Pigot, A. L. (2020). The projected timing of abrupt ecological disruption from climate change. *Nature*, 580(7804):496–501.

Vicente-Serrano, S. M., Beguería, S., and López-Moreno, J. I. (2010). A multiscalar drought index sensitive to global warming: The standardized precipitation evapotranspiration index. *J. Clim.*, 23(7):1696–1718.

Wang, H., Wang, H., Ge, Q., and Dai, J. (2020). The interactive effects of chilling, photoperiod, and forcing temperature on flowering phenology of temperate woody plants. *Frontiers in Plant Science*, 11.

Wessel, P. and Smith, W. H. F. (1996). A global, self-consistent, hierarchical, high-resolution shoreline database. *Journal of Geophysical Research: Solid Earth*, 101(B4):8741–8743.

Zelinka, M. D., Myers, T. A., McCoy, D. T., Po-Chedley, S., Caldwell, P. M., Ceppi, P., Klein, S. A., and Taylor, K. E. (2020). Causes of Higher Climate Sensitivity in CMIP6 Models. *Geophysical Research Letters*, 47(1):e2019GL085782.

Zurell, D., Albert, C. H., Bocedi, G., Briscoe, N. J., Buckley, L. B., Gascoigne, S. J. L., Gonzalez, A., Guillera-Arroita, G., Isaac, N. J. B., Karger, D. N., Lundquist, C. J., Merow, C., Cabral, J. S., Schifferle, K., Velazco, S. J. E., and Urban, M. C. (2026). Biodiversity science and policy need more model intercomparisons. *Nature Reviews Biodiversity*, 2(3):204–215.
